# A Boundary Element Method of Bidomain Modeling for Predicting Cellular Responses to Electromagnetic Fields

**DOI:** 10.1101/2023.12.15.571917

**Authors:** David M. Czerwonky, Aman S. Aberra, Luis J. Gomez

## Abstract

**Objective:** Commonly used cable equation-based approaches for determining the effects of electromagnetic fields on excitable cells make several simplifying assumptions that could limit their predictive power. Bidomain or “whole” finite element methods have been developed to fully couple cells and electric fields for more realistic neuron modeling. Here, we introduce a novel bidomain integral equation designed for determining the full electromagnetic coupling between stimulation devices and the intracellular, membrane, and extracellular regions of neurons.

**Methods:** Our proposed boundary element formulation offers a solution to an integral equation that connects the device, tissue inhomogeneity, and cell membrane-induced E-fields. We solve this integral equation using first-order nodal elements and an unconditionally stable Crank-Nicholson time-stepping scheme. To validate and demonstrate our approach, we simulated cylindrical Hodgkin-Huxley axons and spherical cells in multiple brain stimulation scenarios.

**Main Results:** Comparison studies show that a boundary element approach produces accurate results for both electric and magnetic stimulation. Unlike bidomain finite element methods, the bidomain boundary element method does not require volume meshes containing features at multiple scales. As a result, modeling cells, or tightly packed populations of cells, with microscale features embedded in a macroscale head model, is made computationally tractable, and the relative placement of devices and cells can be varied without the need to generate a new mesh.

**Significance:** Device-induced electromagnetic fields are commonly used to modulate brain activity for research and therapeutic applications. Bidomain solvers allow for the full incorporation of realistic cell geometries, device E-fields, and neuron populations. Thus, multi-cell studies of advanced neuronal mechanisms would greatly benefit from the development of fast-bidomain solvers to ensure scalability and the practical execution of neural network simulations with realistic neuron morphologies.

## 1. Introduction

Device-induced electromagnetic fields are commonly used to treat psychiatric and neurological disorders and study the human brain [1, 2, 3]. Examples of such devices are surface electrodes for transcranial electric stimulation (TES), magnetic coils for transcranial electric stimulation (TMS), and surgically implanted electrodes for deep brain stimulation (DBS). TES has been shown to assist with the treatment of psychiatric disorders and neuro-recovery [1]. TMS was approved by the U.S. Food & Drug Administration (FDA) for treating depression in 2008, obsessive-compulsive disorder in 2018, and smoking addiction in 2020. DBS has been FDA-approved since 1997 for the treatment of tremors due to Parkinson’s disease.

Improving the efficacy of these devices using clinical trials alone to test new protocols and device designs is a costly and time-consuming process. A valuable and complementary approach is to leverage simulations to design devices and stimulation protocols based on biophysical principles governing the interactions between electromagnetic fields and neural activity.

The current standard approach for modeling the neural response to electric fields (E-fields) involves importing E-field dosimetry results into a cable equation solver, such as NEURON [4], to represent extracellular potentials [5, 6] (or an activation function [7, 8]). This approach enables simulations to predict the response of individual neurons to device E-fields [9, 10, 11, 12, 13, 14]. However, this approach comes with inherent assumptions that limit its predictive power. These assumptions include considering cell morphology as filamentary (one-dimensional), assuming that the device-induced E-field remains unaffected by the presence of the cell or its membrane currents, and assuming that the cell is surrounded by an empty conductive extracellular space. The filamentary assumption implies that the neuron’s cross-section is isopotential and that transverse E-fields have negligible effects [15]. In reality, action potentials can be initiated using transverse E-fields, indicating that the cell’s cross-section can result in cell activation [16]. Consequently, correction terms based on passive cell analysis [17] and canonical geometries have been incorporated to enhance the traditional cable equation model [11].

Furthermore, cells are known to significantly disrupt the device E-fields on a local scale [17, 18]. In the human brain, cells are densely packed like other primate brains [19], with only an estimated 15% to 30% of the normal adult brain consisting of extracellular space [20, 21]. The presence of nearby cells can profoundly affect the stimulation that each cell experiences [22, 23, 24, 25].

To address all these limitations for general geometries, finite element method (FEM) bidomain models, also known as “whole” FEM [26, 27], have been proposed by Ying and Henriquez [28] and improved by Neef and Toro [29]. Similar to cable models, this method treats cell membranes as equivalent circuits, creating a bidomain configuration that separates the intra- and extracellular spaces [30, 14, 31]. However, unlike cable-based approaches, this method enables full coupling between cells and device-induced E-fields.

One limitation of this approach is that it requires dense volume meshes that span multiple scales to solve more realistic scenarios, making them computationally intractable for many applications [22, 23].

We propose a boundary integral equation-based bidomain approach that addresses the challenges associated with multi-scale volumetric meshing. Specifically, we solve the boundary integral equation for the charges on each tissue interface using the integral equations of [32] and [33]. For active membranes, we introduce an additional term to model the contribution of transmembrane currents and membrane potentials [34]. These boundary integral equations are solved at each time step, allowing for the incorporation of time-varying membrane properties via auxiliary gating equations. To validate this novel equation, we compare its results with those obtained using our in-house implementation of the FEM method proposed in [29] and the standard cable equation approach.

Additionally, we present a wide range of results to demonstrate the potential applications of our integral equation approach. These include various canonical test cases involving multiple axons, local field potentials (extracellular potentials), cell packing, transverse polarization, DBS electrodes at varying proximity to axons of varying dimensions, and TMS using a spherical head model.

## 2. Methods

### 2.1. Modeling device induced E-fields

The head is divided into nested regions with conductivities denoted from inner to outer as *σ*_1_, *σ*_2_, …, *σ*_*N*_ [Figure 1A]. Each region is separated by boundaries denoted from inner to outer as Γ_1_, Γ_2_, …, Γ_*N*_, each having an outward-pointing normal 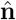. (Note: the results remain valid for non-nested regions, but we assume they are nested to simplify notation.)

**Figure 1:**
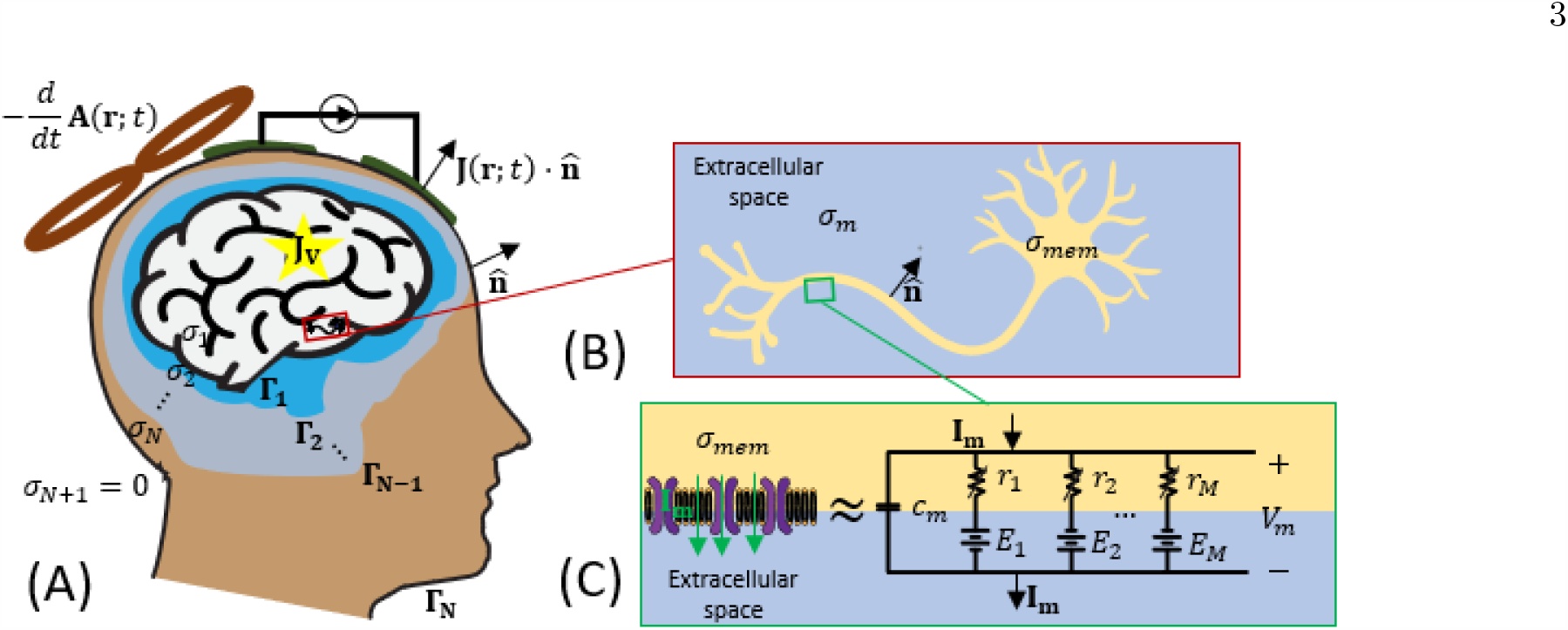
(A) Illustration of the head stimulated by a variety of electromagnetic brain stimulation devices, (B) the neuron cell in the brain, and (C) the neuron membrane and its equivalent circuit model.

Different types of devices induce conduction currents in the head through various mechanisms. Scalp and implanted electrodes are modeled as boundary current injections 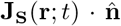, while TMS coils are modeled as volume conduction currents 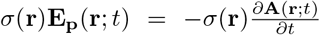 inside the head. The volumetric sources **J**_*V*_ (**r**; *t*) are not used to model device-induced E-fields but are included for generality. In general, these sources will generate a scalar potential *ϕ*(**r**; *t*) that satisfies the following equation and boundary condition:

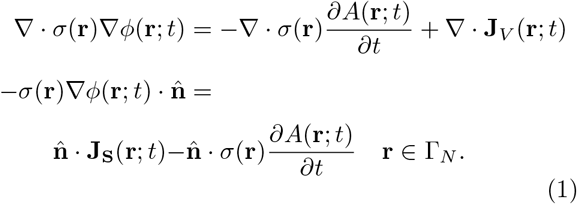

The scalar potential *ϕ*(**r**; *t*) is continuous across any tissue or membrane interface and can be determined from surface charges *ρ*(**r**; *t*) as:

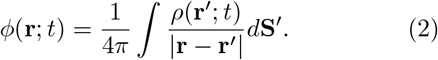

In the appendix, Green’s second identity is applied at each boundary to derive an adjoint integral equation [32] to solve for the charge:

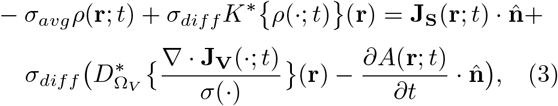

where

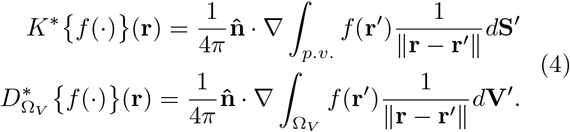

In these equations 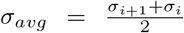 is the average conductivity across the tissue boundary, *σ*_*diff*_ = *σ*_*i*+1_ − *σ*_*i*_ is the difference in conductivity between the outside and inside of the tissue boundary. Note that here we assume that *σ*_*N*+1_ = 0 because it is assumed that the media surrounding the head is insulating. Additionally, 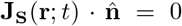 everywhere except at the electrode cathodes and anodes, and Ω_*V*_ is the spatial support of the volume current. The adjoint double-layer operator has been extensively studied and has been shown to accurately predict E-fields for both magnetic stimulation [32, 35, 36, 37] and electric stimulation [32, 36]. The following section will extend the adjoint double-layer potential to include an active membrane. (Note that for notational brevity here and in what follows we have suppressed the spatial dependence of *σ*_*avg*_, *σ*_*diff*_, and 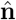 as their values are implied by the boundary location.)

### 2.2. Equivalent circuit neuron membrane models

Membranes that separate the intra- and extracellular space are modeled using equivalent circuits. Equivalent circuits as shown in [Figure. 1 C] have been utilized for over 70 years and are extensively validated by experimental research [38, 39, 40, 41]. The standard formalism for modeling neuron membranes is as follows:

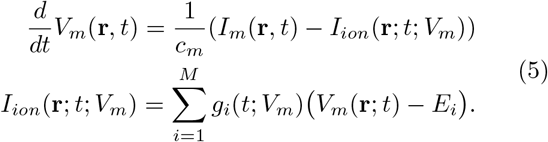

In these equations *I*_*m*_(**r**, *t*) and *V*_*m*_(**r**, *t*) are the transmembrane current and voltage, respectively, *c*_*m*_ is the membrane capacitance, *g*_*i*_ and *E*_*i*_ are the conductance and reversal potential for the ith ion channel, respectively, and *M* distinct ion channels are assumed. In this paper, we aim to introduce the boundary equations and restrict ourselves to Hodgkin Huxley (HH) squid axon model [38]

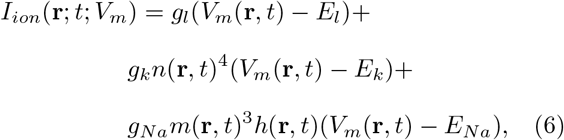

where

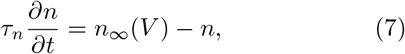

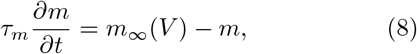

and

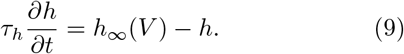

Parameters for the auxiliary gating equations (7,8,9) are given in table 1 of Appendix A. The parameters for (6) are given at the end of section 2.5.

### 2.3. Coupling neuronal membranes to the adjoint double layer potential

The cell is assumed to be in the *m*-th compartment of the head model [Fig. 1A-B]. There will be a jump in the scalar potential across the cell membrane (i.e., between the inside and outside of the cell) equal to *V*_*m*_(**r**; *t*). This jump in the scalar potential is not accounted for in (1) because in its derivation it is assumed that the scalar potential is continuous. To include an active membrane, an additional term must be added to (1) that accounts for the jump in the potential at the cell boundary Γ_*mem*_. This results in the following boundary integral equation, which we call a bidomain integral equation,

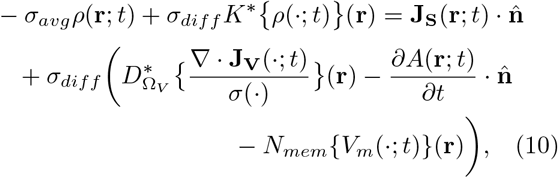

where

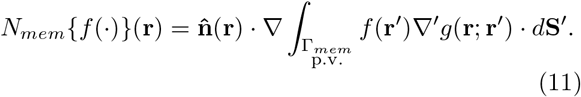

This equation is valid on all interfaces including Γ_*mem*_. Furthermore, the transmembrane current will be equal to the total current flowing out of the cell, which is related to the charge on the surface of the cell as

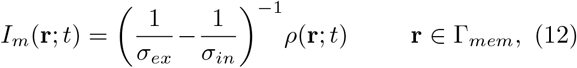

where, *σ*_*ex*_ and *σ*_*in*_ are the extracellular and intracellular conductivities.

### 2.4. Temporal approximation of the membrane equation

Starting from known membrane potential and trans-membrane and ion currents at an initial state *t* = *t*_0_, we solve for *I*_*m*_(**r**, *t*_0_+*n*Δ*t*), *I*_*ion*_(**r**, *t*_0_+*n*Δ*t*), *V*_*m*_(**r**, *t*_0_+ *n*Δ*t*) and *ρ*(**r**, *t*_0_ + *n*Δ*t*), where *n* = {1, 2, …, *N*_*t*_} and Δ*t* is the time step. This is achieved by using the Crank-Nicolson method [42], as employed in [29]. The Crank-Nicolson method is well-known for its unconditional stability when applied to the cable equation [43] and equation (14). Specifically, a first-order central difference method is used for the state equation of the membrane, resulting in the following

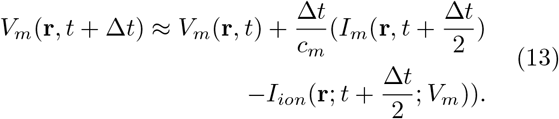

To solve (13), the ionic currents at the half-time step are approximated as the ionic currents at the previous time step, and the membrane currents at the half-time step are approximated using the average of the currents at the next and previous time steps. These approximations result in the following equation:

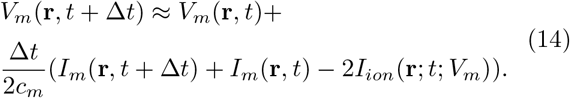

Equation (14) relates *V*_*m*_(**r**, *t* + Δ*t*) to known ionic and membrane currents at the previous time step and *I*_*m*_(**r**, *t* + Δ*t*). Additionally, *I*_*m*_(**r**, *t* + Δ*t*) is related to *ρ*(**r**, *t* + Δ*t*) via (12). These relations for transmembrane current (12) and transmembrane voltage (14) allow us to rewrite our bidomain integral equation in the following form

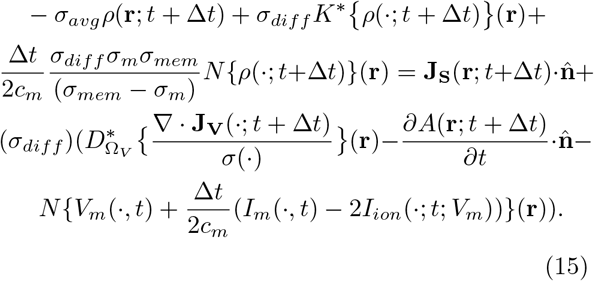

Using known quantities at the previous time step and known sources, equation (15) is solved numerically using the boundary element method to determine *ρ*(**r**; *t* + Δ*t*). Then, *I*_*m*_(**r**, *t* + Δ*t*) and *V*_*m*_(**r**, *t* + Δ*t*) are determined using (12) and (14), respectively. Finally, *V*_*m*_(**r**, *t* +Δ*t*) is used to update the time-dependent ion channel conductances and determine *I*_*ion*_(**r**, *t*+Δ*t*; *V*_*m*_) using (5).

One of the strengths of the time-stepping scheme used here is that the update equations for ionic currents and channel conductance remain unchanged relative to point-wise neuron solvers. Here, we use a backward difference Euler time-stepping scheme for the ionic currents as provided in [40]. First, *V*_*m*_(**r**, *t* + Δ*t*) is used to update the ratio of open and closed ionic gates using the following formula:

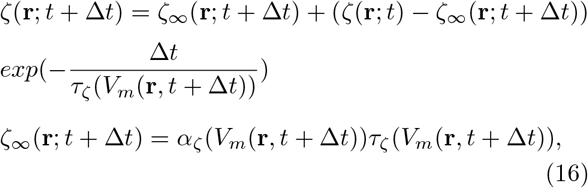

where *ζ* ∈ *n, m, h*. The updated ion gating probabilities are used to determine updated ionic currents as

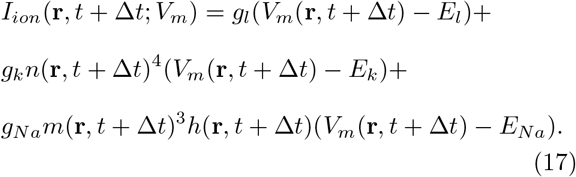

### 2.5. Boundary element methods (BEMs) for approximating the update equation

To approximate the integral equation (15), each boundary is discretized as a mesh consisting of triangles. For the standard adjoint double layer equation, piece-wise constant charges on each triangle are sufficient to achieve accurate results for both the charges and electric fields [35]. However, for the bidomain integral equation, we have an additional hyper-singular operator *N*_*mem*_{*f* (·)} (**r**), which requires a continuous function as input [44]. To meet this requirement, the lowest-order basis functions that enable an accurate approximation of the membrane bidomain equation are linear elements. Consequently, the charges are approximated as linear nodal elements on each node of the triangle mesh.

There was a previous successful attempt for approximating the hyper-singular operator using piece-wise constant functions for studying passive cells [34]. This approach involved using Eq.(6.17) shown in [44] to set the diagonal entries of **N**_**mem**_ to the negative value of all off-diagonal equation coefficients summed together. This regularization method is equivalent to using the representation *N*_*mem*_ {*f* (·) }(**r**) = *N*_*mem*_{ *f* (·) − *f* (**r**)} (**r**) which is only valid when *f* (**r**) is continuous [44]. Although this approach is not mathematically correct, the above regularization appears to produce numerically correct approximations of the hypersingular operator using piece-wise constant basis functions.

The application of a Galerkin procedure to (10) results in the following equation

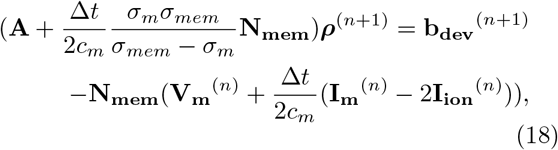

where

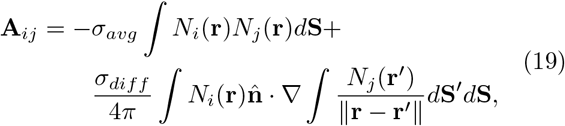

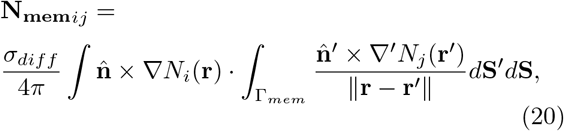

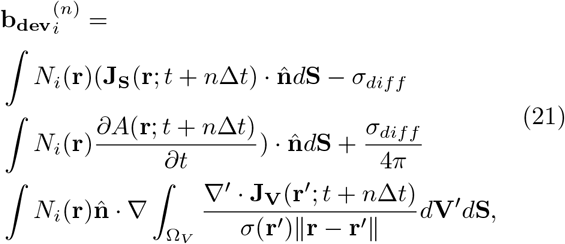

and

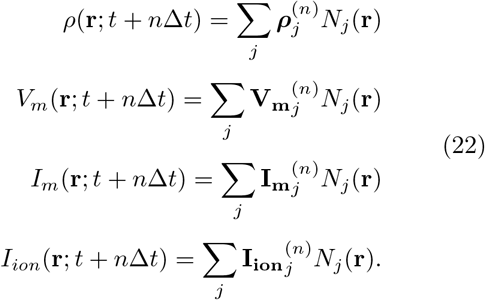

Here, *N*_*i*_(**r**) are nodal elements. The matrix **A** and the forcing vector **b**_**dev**_ correspond to the standard adjoint double-layer equation. The coupling with the active membrane involves adding a single integral operator **N**_**mem**_. This makes the incorporation of membrane equations into existing BEM solvers relatively straightforward.

We explored two approaches to solve (18) for the charges. One approach is to invert the system of equations. Another approach is to solve it iteratively to a relative residual error tolerance of 10^*−*7^ using fast multipole method (FMM) acceleration for the matrix-vector multiplication. Each simulation required over 10 thousand time-steps as a result, the iterative solution using FMM was too slow and the results are not reported here. For all simulation results, the solver is assumed to start at rest with *V*_*m*_(**r**; 0) = −65 mV. We used membrane capacitance*c*_*m*_ = 1*μF/cm*^2^, leak conductance *g*_*l*_ = 3*mS/cm*^2^, peak potassium condutance *g*_*k*_ = 360*mS/cm*^2^, and peak sodium conductance *g*_*Na*_ = 1200*mS/cm*^2^. The reversal potentials were*E*_*l*_ = −54.4 mV, *E*_*k*_ = −77 mV, *E*_*Na*_ = 50 mV.

### 2.6. Summary of Solver Algorithm

As depicted in Fig. 2, each simulation begins with the assumption that the initial state of the cell is known. Subsequently, we solve (18) to calculate the charge at the next time step for every boundary. Using the charge information, we update the transmembrane current *I*_*m*_ for the next time step according to (12). The membrane voltage *V*_*m*_ is then updated via (14). Following this, the membrane voltage is utilized to update the ion channel conductance using the gating equations (16). With the ion channel conductances determined, we calculate the ionic currents for the next time step using (17). Steps one through five are iteratively repeated until the values at all time steps have been determined.

**Figure 2:**
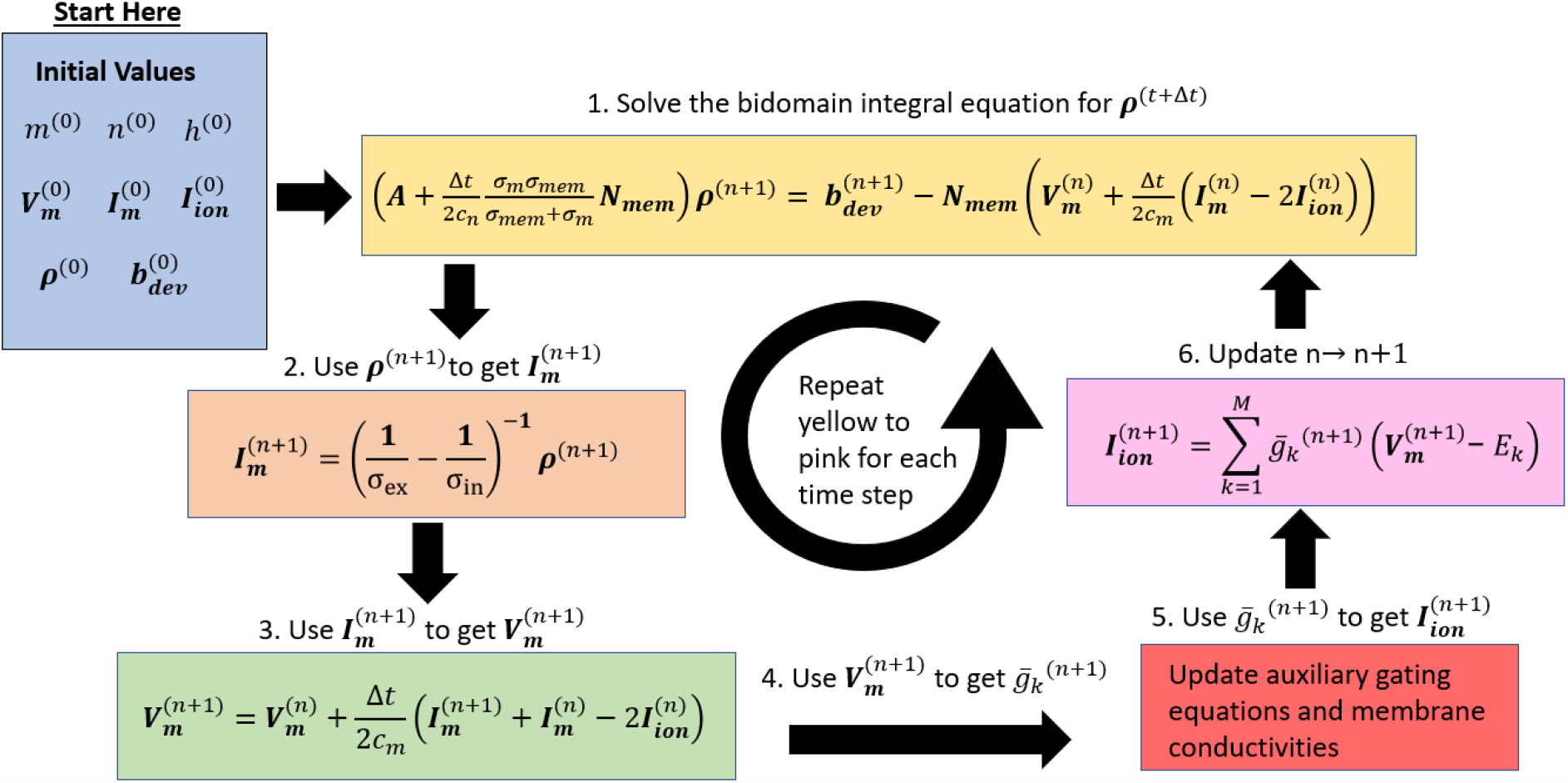
A procedural flowchart of our bidomain BEM algorithm. The procedure is repeated until the charge and membrane information are calculated for all time steps.

### 2.7. Computer simulations

To validate the proposed integral equation solver, we employed the approach described in [27], where extra-cellular voltages are computed from a finite element method (FEM) to drive a cable neuron simulation. The bidomain FEM approach is implemented in-house, based on the description in [29]. For certain aspects of the simulation, we utilized the NEURON simulation environment v8.8.2 [4]. All tasks related to mesh generation, differential equation solutions, and analysis were performed using MATLAB (R2020a, MathWorks, Inc., Natick, MA, USA). The computational resources used for these simulations were provided by the Purdue Bell Cluster, which consists of 5 Rome CPUs running at 2.0 GHz, with a total RAM memory of 10 GB available for each axon in the simulation.

### 2.8. Simulation scenarios & device description

Unless stated otherwise, a time step of 1 *μ*s is used for all simulations. Furthermore, we consider straight axons with a length of 4 mm and a cylindrical cross-section with a diameter of 2 μm. Furthermore, the intracellular and extracellular conductivity are set to 1 S/m and 2 S/m, respectively. This is to match cytoplasm conductivity for a glia cell, which is estimated to be in the range of 0.3-1 S/m [45].

In the single-axon scenarios, all electrodes and coils are driven at threshold current intensities, which we determined empirically with a 1.0 % tolerance using a bisection method.

In the TES scenarios, the extracellular space is represented as a homogeneous hexahedral bath with dimensions of 16 mm × 8 mm × 16 mm as shown by Figure 3(A). The axon is positioned with its midpoint at the center of the hexahedral bath. The electrodes are modeled as having a uniform current distribution on their cross-section. The surface electrodes cover an entire face of the hexahedral bath for the anode and cathode, respectively.

**Figure 3:**
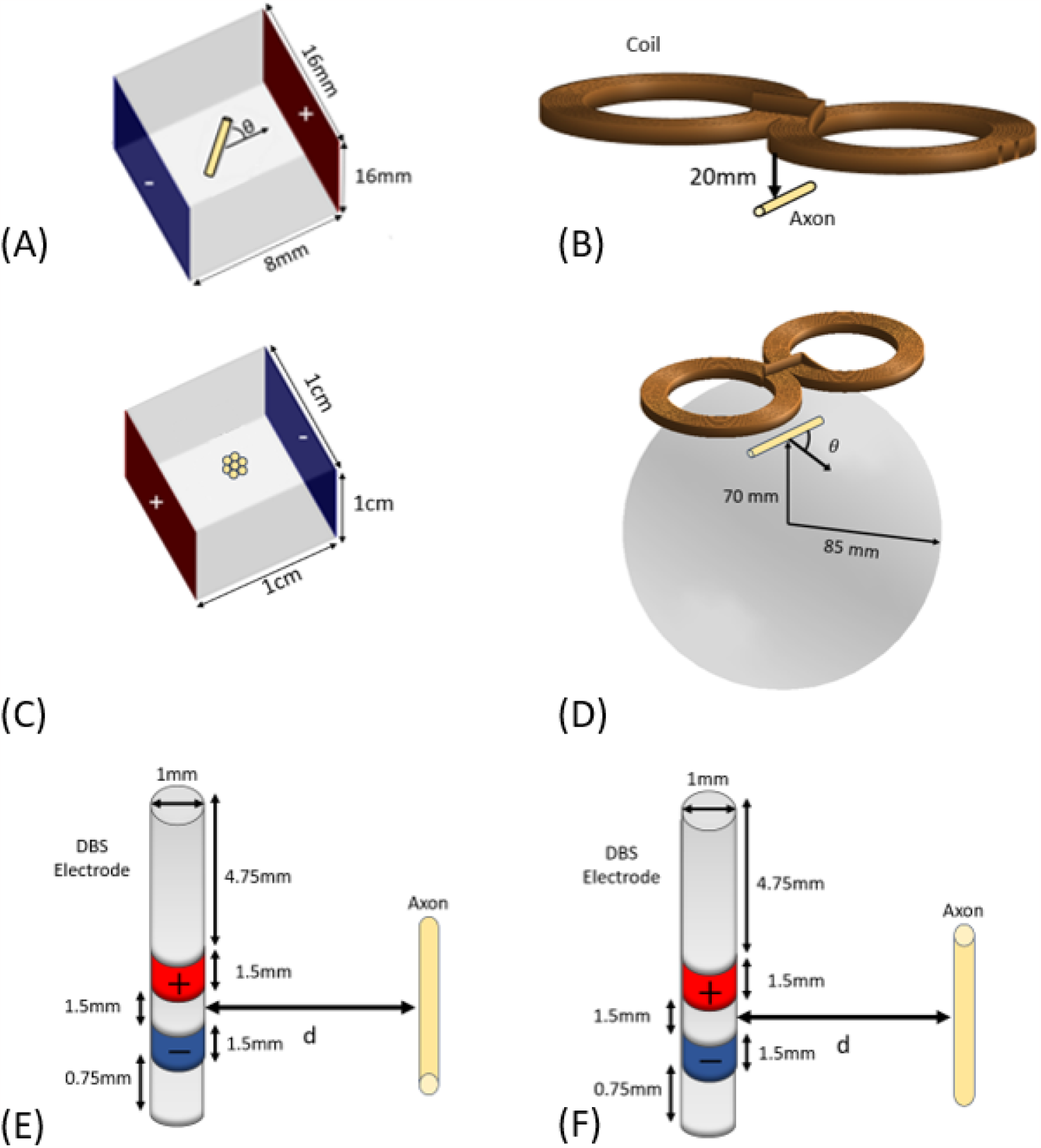
(A) Schematic for TES of an HH axon in a rectangular bath with two surface electrodes denoted by the anode (blue) and cathode (red) stimulating the axon. (B) Schematic for TMS of an HH axon in an unbounded extracellular medium. (C) Schematic of TES of a hexagonal cluster of seven 20 *μ*m diameter spherical cells with 1 *μ*m spacing between one another. (C) Schematic for TMS of an HH axon embedded in a spherical head model. (E) Schematic for DBS perpendicular to an HH axon in an unbounded extracellular medium at various distances *d* of separation between the electrode and an axon. (F) Schematic for DBS parallel to an HH axon in an unbounded extracellular medium at various distances *d* of separation between the electrode and an axon.

For Figure 3(B), we consider an unbounded extracellular space for the longitudinal comparison scenario. For Figure 3(D), we consider an 85 mm radius spherical head. An 85 mm radius spherical head is a common choice of simple head model in TMS simulation studies [46, 47, 48].

An 85 mm figure-eight coil is placed 90 mm above the center of the spherical head. The axon is placed in a variety of orientations with its midpoint 70 mm above the center of the spherical head. The placement of the neuron with respect to the figure-eight coil falls within the range used in studies of neuron stimulation by figure-eight coils [12, 49]. The coil model consists of 24,426 electric dipoles with inductance of 17.061 *μ*H, and resistance of 6.3 mΩ. The underdamped pulse waveform shape is determined assuming the coil is driven by a 185 *μ*F charged capacitor.

For the DBS scenarios, we ran simulations of a cylindrical DBS electrode in an unbounded extracellular space at a distance *d*, ranging from 50 nm to 1 cm away from an axon. The axon is placed in two orientations, perpendicular to the electrode or parallel to the electrode. The electrode is represented as a cylindrical rod with a radius of 0.5 mm and a length of 10 mm, as illustrated in Figure 3 (E) and (F). The anode is modeled as a 1.5 mm thick band located 0.75 mm from one end of the electrode, while the cathode is a 1.5 mm thick band positioned 1.5 mm above the anode. The DBS electrode is assumed to have uniform current distribution in the cathode and anode regions, and the inside of the electrode is considered insulating. We drive the electrode with a square wave pulse that starts at 0.2 ms and lasts for 5 ms. The lengthwise midpoint of the axon is aligned with the midpoint between the electrode anode and cathode in all DBS scenarios.

Additionally, we consider DBS simulations with a longer and a thicker axons. We set our long axon to have dimensions of 10 mm in length and 2 *μ*m in diameter. The thick axon is 4 mm in length and 40 *μ*m in diameter.

For the extracellular field calculation scenarios, we place an HH axon with its center at the origin of an unbounded conductive extracellular space and connect a current source to the ends of the axon. These currents are chosen at the activation threshold level. Then, we calculate the extracellular potentials nearby (*<* 200 *μ*m) of the space outside the axon membrane using the relation (B.9) of Appendix B. The extracellular fields scale in an inversely proportional fashion to the conductivity [50, 51]. We chose the extracellular conductivity to be 0.1 S/m for these simulation scenarios. The value of 0.1 S/m is low enough to get strong extracellular fields but remains realistic. Conductivity estimates of the extracellular space around myelinated cells are in the range of 0.28 S/m to 5.5 S/m [52].

To investigate how the distortion of the local E-field may affect nearby cells, we consider a bath containing a hexagonal cluster of seven 20 *μ*m diameter spherical cells with 1 *μ*m spacing and arranged as shown in Figure 3 (C). Four distinct types of scenarios are considered: (1) only one cell is present, (2) all of the cells are active, (3) only one cell is active and others have a conductivity of 2 S/m, (4) only one cell is active and others are insulating (i.e. initial polarization assumption [17]). For each of the four types of scenarios, we determine the activation threshold of each cell.

### 2.9. Mesh construction

We generated meshes for these physical models using functions from the Geometry Processing Toolbox (gptoolbox) in MATLAB. For all axons, we created meshes with a minimum of 810 nodes (8 nodes around the circumference and 100 compartments lengthwise) and 1,616 triangles using the cylinder mesh function. High-resolution axon simulations, consisting of 3,210 nodes and 6,416 triangles, were also performed to evaluate solution convergence, and we determined that higher resolution is unnecessary.

For modeling the extracellular space in TES, we employed the surface mesh generated by the cube function from gpttoolbox, with a resolution setting of 20 (resulting in 2,168 nodes and 4,332 triangles) to represent the hexahedral bath. This mesh is adjusted to match the specified dimensions of the hexahedral bath mentioned earlier.

In the case of the spherical head mesh, we define a nearly uniform spherical mesh by barycentrically refining an icosahedron mesh three times and projecting the nodes to the sphere surface.

## 3. Results

### 3.1. Longitudinal comparison of bidomain BEM

In this section, we examine the effect of longitudinal fields from TES and TMS on the membrane voltage obtained at two specific points along an axon’s length. We consider membrane voltages at 1*/*5 th and 4*/*5 th of its total length.

For the TES longitudinal comparison scenario, the results are presented in Fig. 4(a). The outcomes obtained from the bidomain FEM (see the supplement, Figure S1), the cable equation (CE), and our proposed bidomain BEM all exhibit close agreement. As expected for a straight axon in a uniform E-field, the action potential initiates at the termination closest to the negative electrode and propagates down the axon.

**Figure 4:**
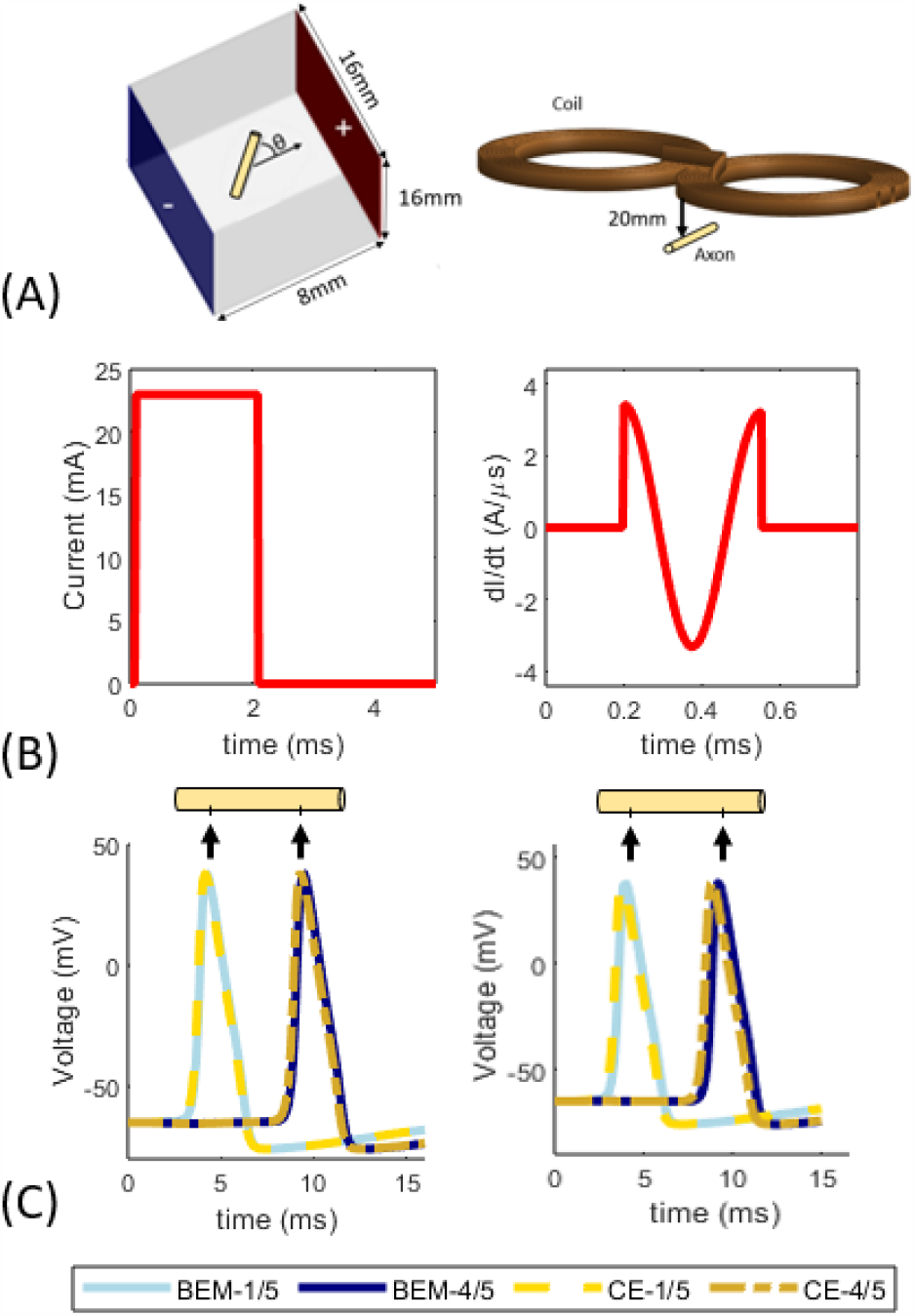
(A) Schematics of the respective TES and TMS longitudinal comparison scenarios. (B) Plots of the waveforms used to drive the electrode and coil devices respectively. (C) Results of TES and TMS by longitudinal stimulation which are validated by the cable equation. The transmembrane voltage is sampled at 1/5th and 4/5th across the length of the axon as denoted by the arrows.

The TMS longitudinal comparison scenario and its corresponding results are displayed in Figure 4(b). Here, we present results using bidomain BEM and the cable equation. Bidomain FEM is excluded in this case due to the computational intractability of creating a volume mesh to represent the open extracellular space. The bidomain BEM and CE exhibit good agreement.

### 3.2. Cell rotation in a uniform E-field

In this section, we investigate the impact of transverse stimulation. Figure 5 (A) presents the results of axons being stimulated by electrodes while oriented at various angles relative to the induced E-field from the electrode. It is well-established that the activation threshold varies with angle [11]. Our results closely align with previously reported trends, showing an increase in the activation threshold as the angle increases. Notably at an angle of 90°, the activation threshold diverges to ∞ for the standard cable equation. This divergence occurs due to the neglect of transverse polarization phenomena in the standard cable equation. However, our bidomain BEM results approach a finite threshold value for transverse stimulation (*θ* = 90°).

**Figure 5:**
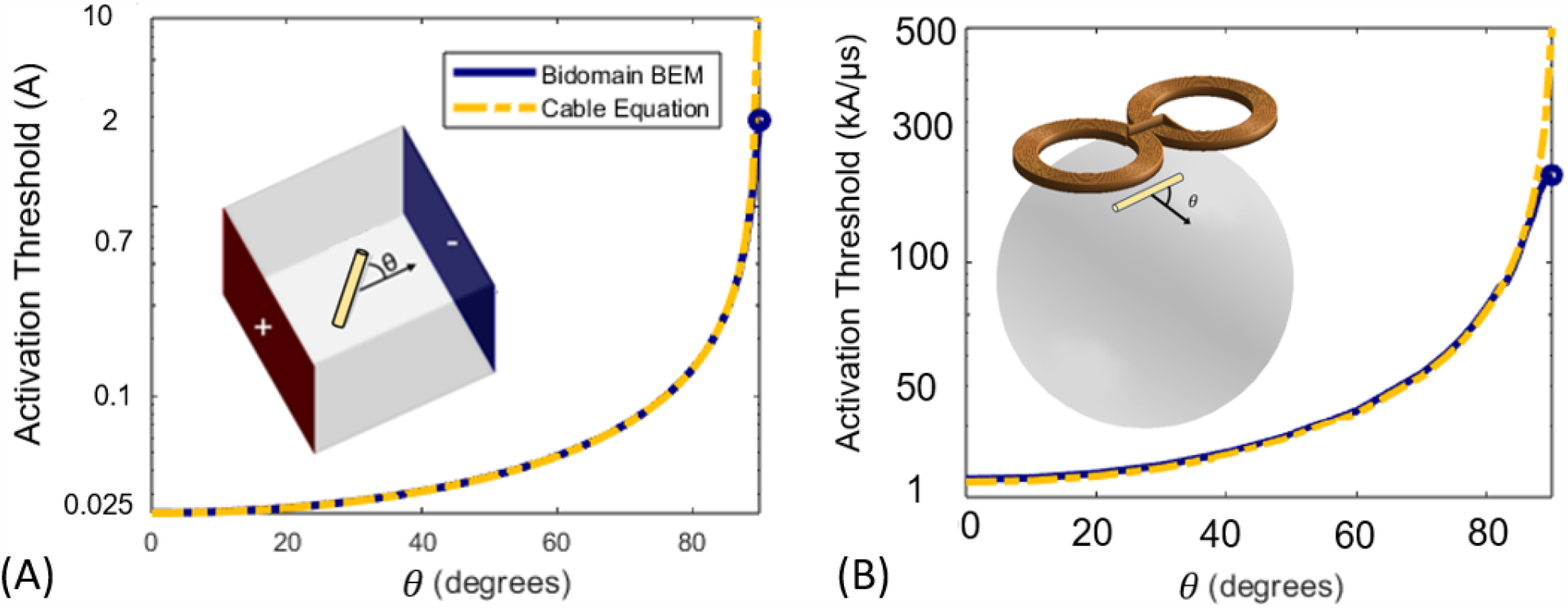
Activation threshold computed by cable equation and bidomain BEM as a function of axon orientation. (A) Thresholds calculated for TES. (B) Thresholds calculated for TMS in a homogeneous spherical head model.

### 3.3. TMS with a spherical head model

Bidomain FEM-based approaches necessitate volume meshes that span multiple orders of magnitude when considering a single cell within a head model.

In contrast, BEM does not impose such meshing requirements, rendering it a more practical choice. Here, we demonstrate that a bidomain BEM approach can be effectively employed for multi-scale simulations where FEM cannot be employed. In Figure 5 (B), we investigate the activation threshold of cells within a spherical head model as a function of cell orientation. Notably, we observe the same trends for TMS as we did for TES. Specifically, the activation threshold increases with the angle. At 90°, the activation threshold diverges to ∞ for the standard cable equation. However, our bidomain BEM results approach a finite threshold value for transverse stimulation (*θ* = 90°).

### 3.4. Deep brain stimulation

Next, we simulated DBS of multiple axon geometries in both parallel and perpendicular orientations as a function of distance from the stimulating electrode. For parallel orientations, the results obtained with bidomain BEM and the cable equation agree with less than 9% difference maximum and less than 2% on average, as illustrated in Figure 6. We attribute this agreement to the fact that the E-field generated by the electrode is predominantly parallel to the cathode and anode, and thus, it aligns mostly parallel to the axon. For the axons that are 10 mm in length, the action potential initiates from the center of the axon. For the axons that are 4 mm in length, the action potential initiates at one of the terminals. The activation threshold remains similar for distances up to 100 *μ*m from the electrode but increases for cells located farther away. In contrast, the perpendicular configuration generates predictions that differ by a minimum of 38% on average. As the distance between axon and electrode grows the difference between the cable equation and bidomain BEM predictions grows rapidly.

**Figure 6:**
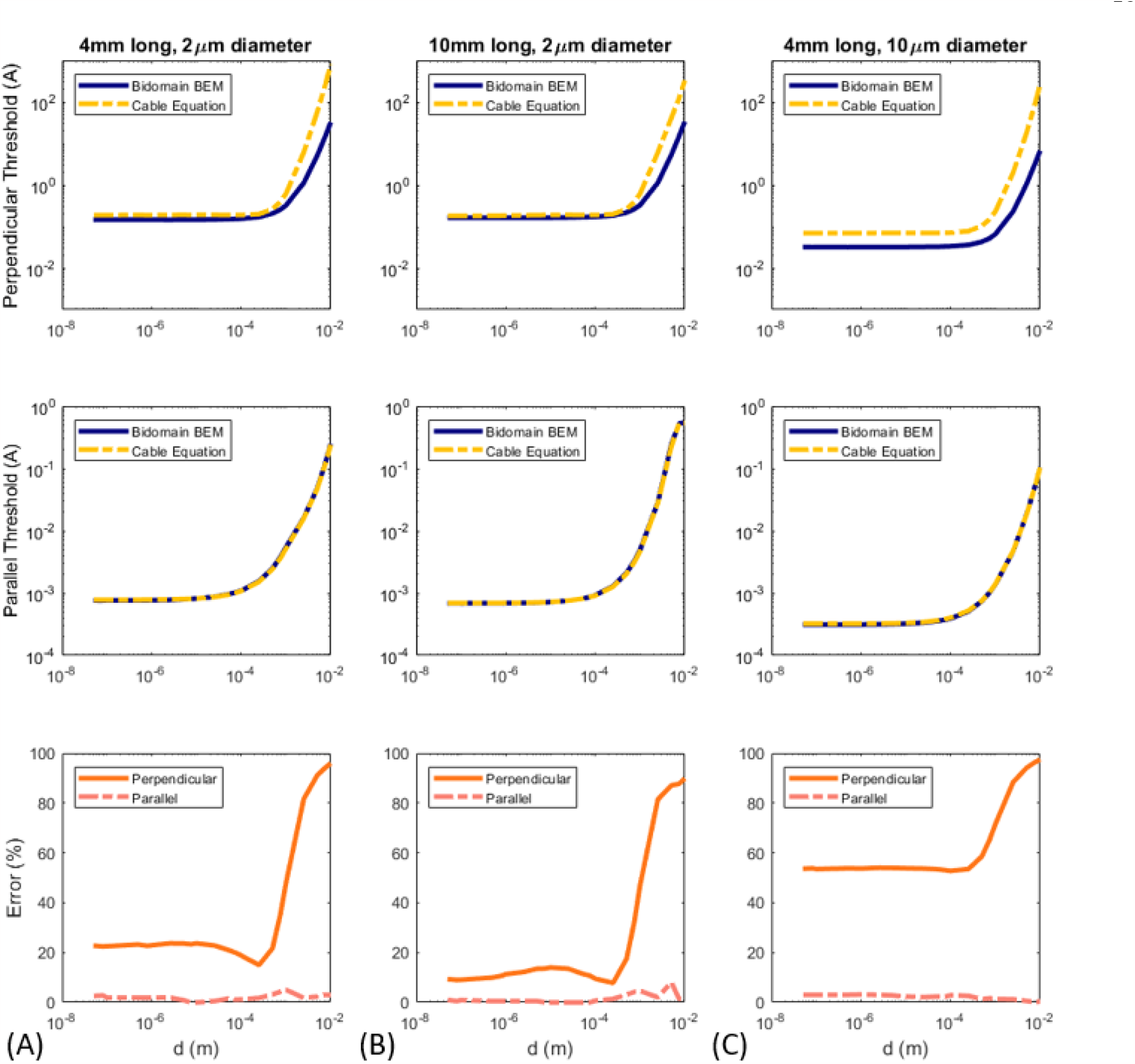
DBS activation threshold as a function of distance *d* to electrode for different axon geometries and orientations. Threhold–distance curves shown for (A) 4mm long, 2*μ*m diameter axon, (B) 10mm long, 2*μ*m diameter axon, and (C) 4mm long, 10*μ*m diameter axon. Within each column, the top and middle row show threshold for the axon perpendicular and parallel to the DBS electrode, respectively, and the bottom row represents the percent error between the bidomain BEM and cable equation with respect to the cable equation.

### 3.5. Modeling extracellular potentials

The bidomain BEM can also be applied to simulate the extracellular potentials generated by neural sources and recorded by external devices. Fig. 7 shows the spatial and temporal profile of the extracellular potentials for an axon stimulated at its activation threshold by a direct current injection. The waveforms replicate the characteristic peaks of extracellular action potential recordings, generated by the capacitive, inward sodium, and outward potassium currents. The peak extracellular potential quickly decays in the transverse 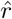 direction, reaching half of the peak strength at r = 10 *μ*m. Additional simulations not provided showed an inversely proportional relationship of extracellular potentials to extracellular conductivity as seen in other works [50, 51].

**Figure 7:**
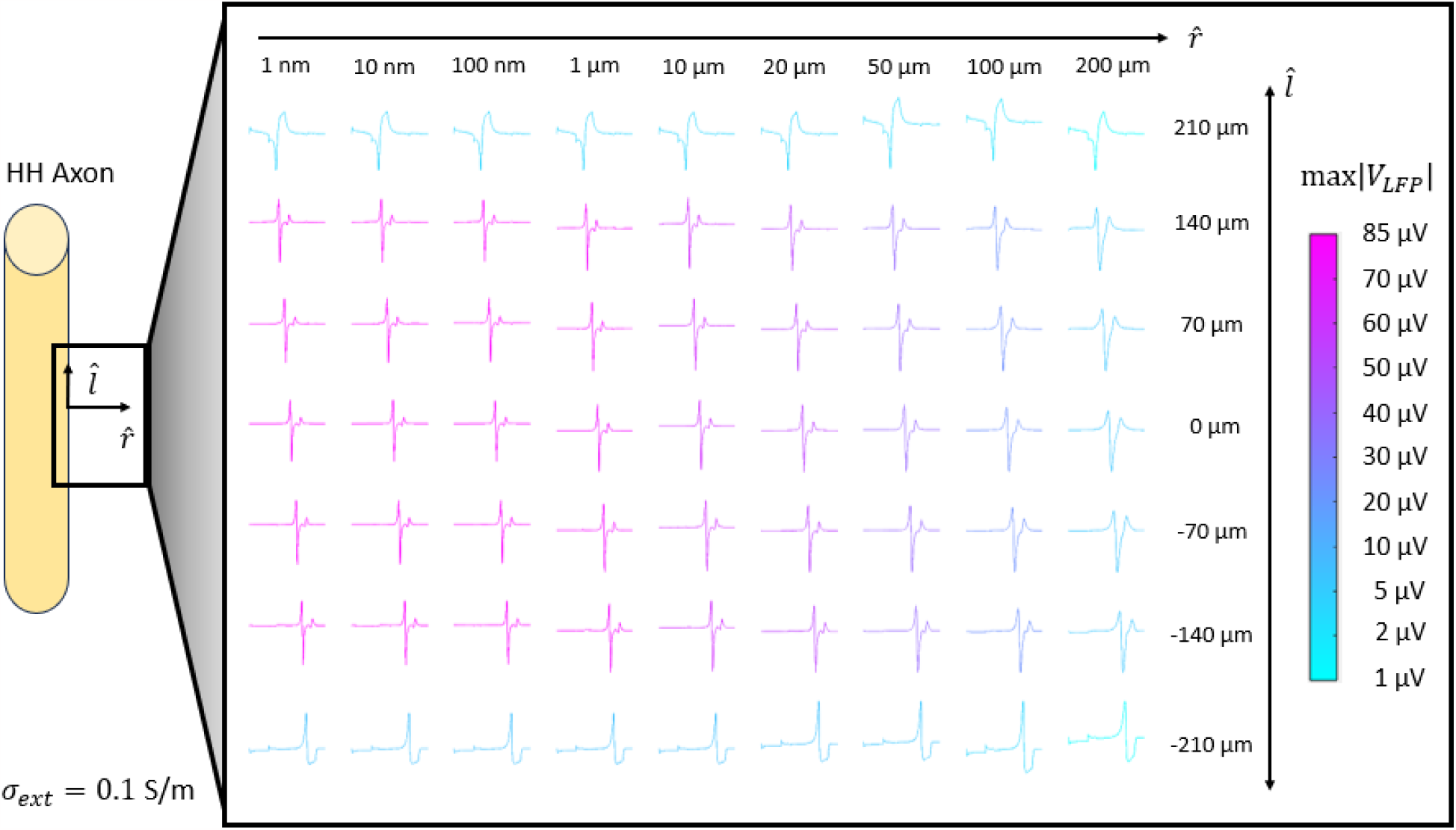
A schematic detailing the region where the data originates is on the left. The plot shows extracellular potential waveforms sampled at the positions on the l-r grid. The colors indicate the magnitude of the largest peak of each respective voltage waveform.

### 3.6. Modeling the effects of neighboring cells

Here, we consider how the presence of multiple cells affects single-cell activation thresholds. To observe these subtle effects, we measured the activation threshold of seven tightly packed spherical cells as shown in Figure 8. In isolation, each spherical cell had the same threshold of 107 mA at their respective positions. Including all seven cells with active membranes increased their activation thresholds depending on their location within the assembly. The center cell threshold was 7.83% higher than the average threshold of the surrounding cells. To determine whether the effects are due to cell activity or shielding we considered one cell active and others as passive conductors (*σ* = 2 S/*m*). We found including adjacent cells with passive membranes also increased activation thresholds relative to that of the cells in isolation; however, the increase was not sufficient to explain the increase in the active case. Finally, we considered one cell active and others as insulators (*σ* = 0 S/*m*), which isolates the state of initial polarization[53]. The resulting thresholds are identical to those when the cells are considered active.

**Figure 8:**
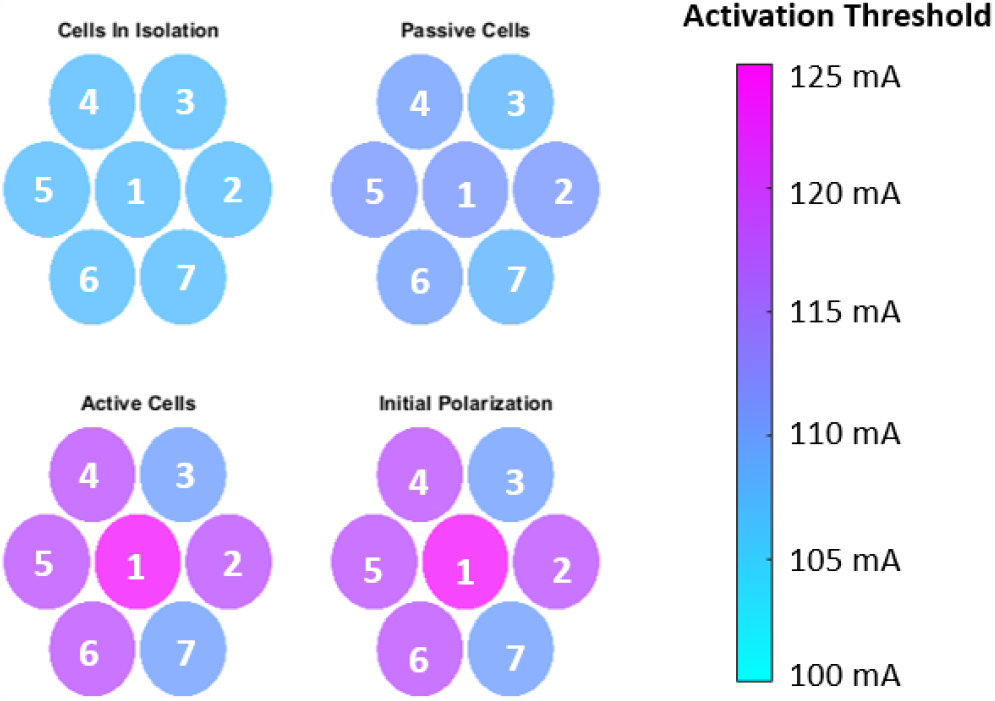
Color plots of the threshold values concerning each spherical cell and the properties of its surrounding cells.

## 4. Discussion

The standard cable equation approach neglects the influence of the cell cross-section, the effect of the cell on the local E-field, and the cell to cell interactions [27, 29, 28]. Bidomain approaches do not require these simplifying assumptions but necessitate additional computational resources. The FEM bidomain approaches require a volume mesh of the cell and surrounding media and capture the exchange of ions across cell membranes by equivalent impedance boundary conditions between neuron cells and the extracellular space. We introduced a BEM-based bidomain solver that avoids the need for volumetric meshing, thereby reducing the computational cost associated with the meshing of bidomain methods. Our longitudinal comparison results indicate that bidomain BEM matches the longitudinal capabilities of the bidomain FEM and the cable methods. Furthermore, the bidomain BEM can be easily applied to scenarios involving large platforms (e.g., head models), and the relative placement of devices to cells can be reconfigured without generating a new mesh.

### 4.1. Assessment of transverse polarization mechanisms

In our TES results (Figures 4(A) and **??** (A)), the cable equation and bidomain BEM approach typically demonstrate good agreement in various scenarios, except for pure transverse stimulation. This pattern also holds for different axon-field orientations in TMS simulations, as illustrated in Figure 5 (B). It’s worth noting that a significant difference between the cable equation and our bidomain BEM only becomes apparent when the angle of the axon with respect to the longitudinal axis exceeds 89°. Our findings indicate that coil currents on the order of 100 kA/*μ*s are necessary for transverse stimulation of an HH axon, consistent with the results observed in Wang et al.’s modified cable equation study [11]. While transverse polarization is impactful in specific cases, it is generally negligible in most practical applications.

These special cases arise when the local E-field exhibits a high magnitude in the transverse direction. For instance, we observe more pronounced disagreements when a DBS electrode is positioned perpendicular to the axon, as illustrated in Figure 6. When the DBS electrode aligns parallel to the axon, we achieve predominantly longitudinal stimulation, resulting in a strong agreement between the cable equation and bidomain BEM. However, when the DBS electrode is oriented perpendicular to the axon, there is limited longitudinal stimulation. Consequently, we observe significant discrepancies between the cable equation and bidomain BEM stimulation thresholds.

In general, our results demonstrate that transverse stimulation necessitates orders of magnitude higher E-field magnitude compared to longitudinal stimulation to elicit an action potential. These findings strongly suggest that longitudinal stimulation is the predominant mechanism for neuron activation via an electrical field.

### 4.2. Modeling of Extracellular Potentials

Our findings match those of [50, 51], which indicate that extracellular potentials are strongest in low-conductivity extracellular spaces. The results with *σ*_*ext*_ = 0.1 S/m most clearly displayed both the extracellular field spatial characteristics. We ran analogous simulations at higher values one of which is included in the supplementary document as Figure S6. We observed that lower extracellular conductivity allowed for higher magnitudes of extracellular potentials, which is consistent with other studies [50, 51, 54, 55].

### 4.3. Modeling a Cluster of Spherical Cells

The results of Figure 8 indicate that shielding of the E-field due to surrounding spherical cells affects activation thresholds strongly. Moreover, we observe that cells in isolation result in a lower threshold than when surrounding cells are considered. For example, when we model a cell in the presence of other cells, we see that the threshold increases up to 15.8% from the threshold in isolation. These results corroborate our claim that the presence of surrounding cells is a relevant determinant of the electric field that cells experience, especially given that neurons are tightly packed.

### 4.4. Significance of the bidomain BEM

For studies with cells in isolation, our results indicate that the conventional (and modified) cable equation is a judicious choice of simulation method. However, when conducting studies on intact neural tissue, a bidomain FEM /BEM or novel cable equation formalism is necessary to capture both the device and neuron-generated fields simultaneously. Generating high-quality volume meshes that span multiple scales can be computationally cumbersome with FEM equivalents. For instance, the use of a spherical head model, often employed as a substitute for an MRI head model, would demand an excessive number of elements to make it suitable for bidomain FEM modeling. In contrast, we have demonstrated that using a BEM approach, TMS simulations, including spherical head models, can be accomplished with ease. Moreover, the flexibility to relocate our device and axon positions without the need to regenerate a mesh makes bidomain BEM a more practical choice, as it enables versatile cell placement without the requirement to generate new volume meshes for each placement.

Bidomain approaches require the solution of a BEM/FEM system of equations and updating of neuron gating unknowns at each time step. Cable equation solvers, in contrast, only require the determination of the E-field once, and only the cable equation must be solved at each time step. As such, the cable equation requires significantly less computational overhead than bidomain methods. Our results indicate that to generate insight and perform single-cell simulations, cable equation-based approaches should be the method of choice. However, for multi-cell scenarios, fast bidomain methods that are scalable and can be executed to model the dynamics of a network of neuron cells with realistic representations of their morphology are necessary. In the context of modeling blood flow by fast BEM solvers have been used to sucessfully model 200 million deformable red blood cells (90 billion unknowns in BEM) [56]. These simulations indicate that modeling a network of neurons is well within our reach with existing computational algorithms and infrastructure.

## 5. Conclusion

We introduced a novel bidomain integral equation for modeling the electric response of neuron cells to device-induced E-fields. Our study includes several canonical test cases, including scenarios with multiple cells, transverse polarization, DBS electrodes at varying proximity to multiple axon geometries, and TMS with a spherical head model. The study results indicate that (1) the cable equation is a sufficient choice for a majority of simulations, (2) longitudinal stimulation serves as the primary activation mechanism for electromagnetic brain stimulation, and (3) multi-cell studies of advanced mechanisms would greatly benefit from fast-bidomain or hybrid cable-BEM solvers to ensure scalability and the practical execution of neural networks simulations with realistic neuron morphologies. Thus, our future efforts will focus on developing fast bidomain solvers to fully incorporate realistic neuron morphologies.

**Table A.1:**
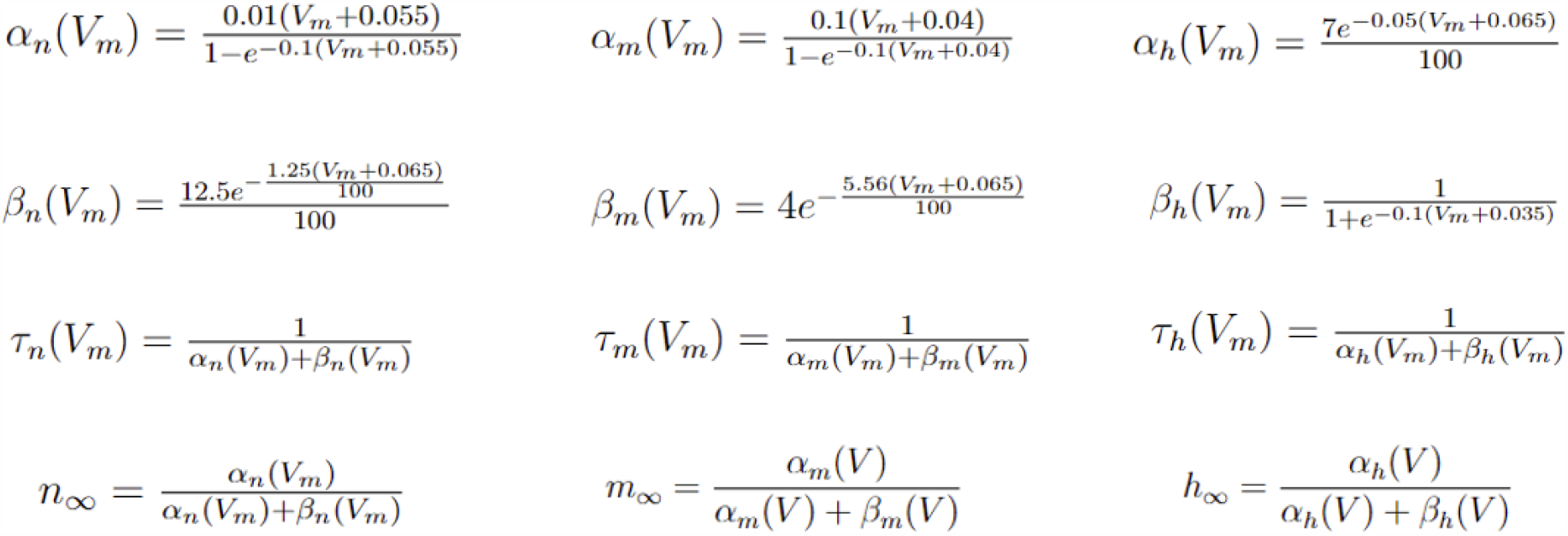
Hodgkin-Huxley axon gating probability and associated auxiliary variable definitions. (7,8,9) [40].

## Supporting information

Bidomain BEM Supplemental

## 6. Acknowledgements

Preliminary results for this work have been presented at the Applied Computational Electromagnetics Society (ACES) 2023 conference (March 2023, Monterey, California, USA)[57], the 2023 IEEE MTT-S International Conference on Numerical Electro-magnetic and Multiphysics Modeling and Optimization (NEMO’2023, June 2023, Winnipeg, Canada), the IEEE International Symposium on Antennas and Propagation Society and USNC-URSI Radio Science Meeting (IEEE-AP-S 2023, July 2023, Portland, Oregon, USA), and the Brain and Human Body Modeling conference 2023 (BHBM 2023, August 2023, Boston, Massachusetts, USA).

Research reported in this publication was supported by the National Institute of Mental Health of the National Institutes of Health under Award Number R00MH120046. The content of the current research is solely the responsibility of the authors and does not necessarily represent the official views of the National Institutes of Health.

## Appendix A

### Hodgkin-Huxley Parameters

Table A.1 shows the rate constants *α*(*V*_*m*_) and *β*(*V*_*m*_), the gating time constants *τ* (*V*_*m*_), and the steady state target values *m*_*∞*_, *n*_*∞*_, *h*_*∞*_. The rate constants must be calculated at each time step to update the gating equations (7,8,9) and the ion current (6).

## Appendix B

### Adjoint Double Layer Equation

In a region Ω_*j*_ with a boundary Γ_*j*_, the scalar potential satisfies the following

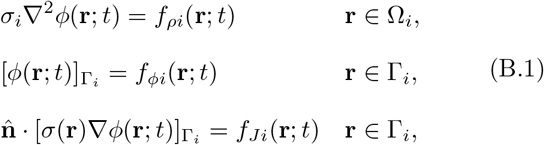

for *i* ∈ {1, …, *N* + 1}. Here we define the jump of a function as 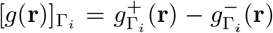, where 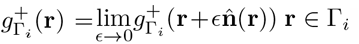 and 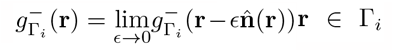 (i.e. the difference between its value just outside and just inside a compartment), and 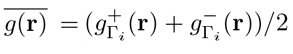 as the average of a function across the boundary. For example, a continuous function, *f* (**r**) will have 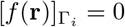, and 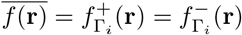, where *i* = 1, ..*N*. The forcing functions *f*_*ρi*_, *f*_*ϕi*_, and *f*_*Ji*_ each model implanted electrodes, potential differences across a thin cell membrane, and electrode and TMS induced charges, respectively. Green’s second applied to a homogenous region is

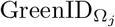 (**r**) :

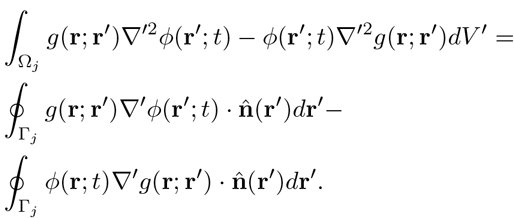

We assume 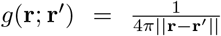 is the Green’s function for the Laplace operator which satisfies ∇^2^*g*(**r**; **r’**^*′*^) = −*δ*(**r** – **r’**^*′*^). This results in

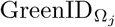 (**r**) :

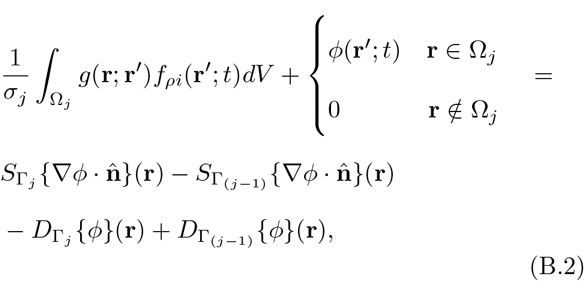

where

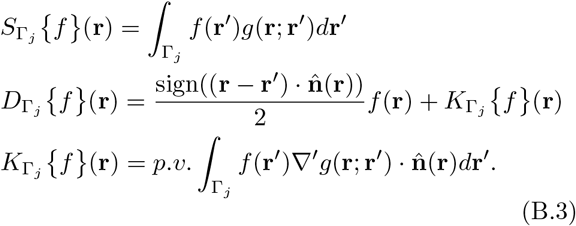

The double layer potential equation is formed by applying 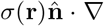 to the Green’s identity above and summing it over all regions

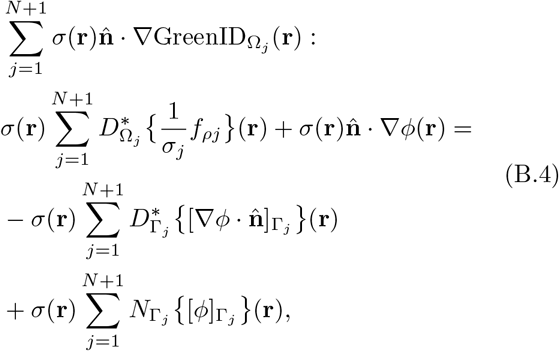

where

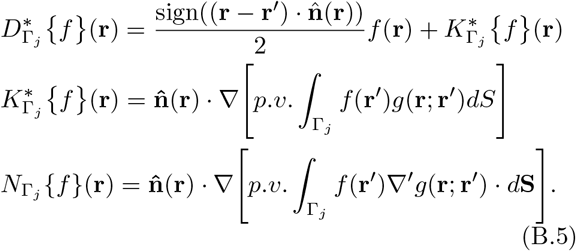

Taking the jump:

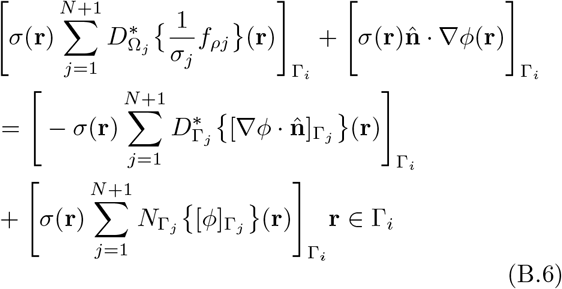

Assuming that the volume sources do not extend across any boundaries and applying the jump to the operators results in the following

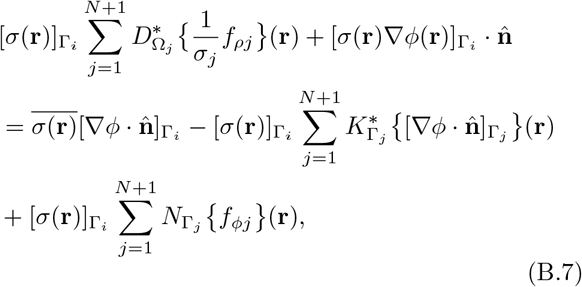

Substituting 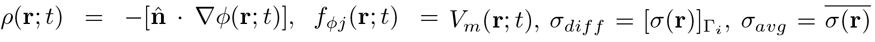 into the above expression and rearranging terms gives the general form as seen in equation 10

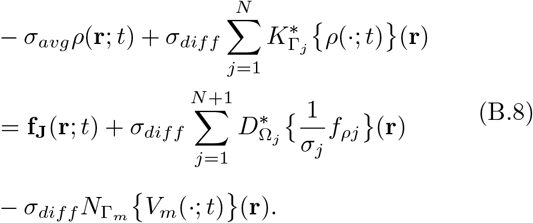

In equation B.8, the hyper-singular operator acting on *V*_*m*_ represents the normal component of the E-field generated by the discontinuity of the scalar potential. With the assumption that *f*_*ρj*_ = 0, the scalar potential can be decomposed into two parts as shown by [58] (Eq. 11) and [33] (Eq. 33):

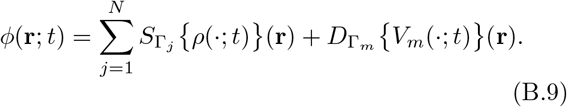

Here the following relation 12 results in the equation given by [58] and can be used to describe the extracellular fields due to transmembrane current and voltage.

## References

[1] Dennis Q. Truong, Niranjan Khadka, and Marom Bikson. Transcranial Electrical Stimulation, pages 271–292. Springer International Publishing, Cham, 2020.

[2] Walter Paulus. Transcranial Magnetic Stimulation. Elsevier Imprint, 2003.

[3] Michael S Okun. Deep-brain stimulation for parkinson’s disease. New England Journal of Medicine, 367(16):1529–1538, 2012.

[4] Michael L Hines and Nicholas T Carnevale. The neuron simulation environment. Neural Comput., 9(6):1179–1209, 1997.

[5] Donald R McNeal. Analysis of a model for excitation of myelinated nerve. IEEE. Trans. Biomed. Eng., BME-23(4):329–337, 1976.

[6] Srikanthan S Nagarajan and Dominique M Durand. A generalized cable equation for magnetic stimulation of axons. IEEE Transactions on Biomedical Engineering, 43(3):304–312, 1996.

[7] Frank Rattay. Analysis of models for external stimulation of axons. IEEE. Trans. Biomed. Eng., BME-33(10):974–977, 1986.

[8] Frank Rattay. Ways to approximate current-distance relations for electrically stimulated fibers. J. Theor. Biol., 125(3):339–349, 1987.

[9] Jose Gomez-Tames, Ilkka Laakso, Takenobu Murakami, Yoshikazu Ugawa, and Akimasa Hirata. Tms activation site estimation using multiscale realistic head models. J. Neural Eng., 17(3):36004–036004, 2020.

[10] Andreas Fellner, Amirreza Heshmat, Paul Werginz, and Frank Rattay. A finite element method framework to model extracellular neural stimulation. J. Neural Eng., 19(2):22001–, 2022.

[11] Boshuo Wang, Aman S Aberra, Warren M Grill, and Angel V Peterchev. Modified cable equation incorporating transverse polarization of neuronal membranes for accurate coupling of electric fields. J. Neural Eng., 15(2):026003, 2018.

[12] Boshuo Wang, Warren M. Grill, and Angel V. Peterchev. Coupling magnetically induced electric fields to neurons: Longitudinal and transverse activation. Biophys. J., 115(1):95–107, 2018.

[13] Bradley J Roth and Peter J Basser. A model of the stimulation of a nerve fiber by electromagnetic induction. IEEE. Trans. Biomed. Eng., 37(6):588–597, 1990.

[14] Peter J Basser and Bradley J Roth. New currents in electrical stimulation of excitable tissues. Annu. Rev. Biomed. Eng., 2:377–397, 2000.

[15] Robert E. Taylor. Cable theory. In Electrophysiological Methods: Physical Techniques in Biological Research, number v. 6, pt. 2 in Physical techniques in biological research, chapter 4, pages 219–262. Elsevier Science, 1963.

[16] Jarmo Ruohonen, Marcela Panizza, Jan Nilsson, Paolo Ravazzani, Ferdinando Grandori, and Gabriella Tognola. Transverse-field activation mechanism in magnetic stimulation of peripheral nerves. Electroencephalography and Clinical Neurophysiology/Electromyography and Motor Control, 101(2):167–174, 1996.

[17] Wanda Krassowska Neu. Analytical solution for time-dependent potentials in a fiber stimulated by an external electrode. Med Biol Eng Comput, 54(11):1719–1725, 2016.

[18] Jaakko Malmivuo and Robert Plonsey. Bioelectromagnetism: Principles and Applications of Bioelectric and Biomagnetic Fields. Oxford University Press, 10 1995.

[19] Suzana Herculano-Houzel. The human brain in numbers: a linearly scaled-up primate brain. Frontiers in human neuroscience, page 31, 2009.

[20] Eva Syková. The extracellular space in the cns: its regulation, volume and geometry in normal and pathological neuronal function. The Neuroscientist, 3(1):28–41, 1997.

[21] Eva Syková and Charles Nicholson. Diffusion in brain extracellular space. Physiological reviews, 88(4):1277–1340, 2008.

[22] Hamish Meffin, Bahman Tahayori, David B Grayden, and Anthony N Burkitt. Modeling extracellular electrical stimulation: I. derivation and interpretation of neurite equations. J. Neural Eng., 9(6):065005, 2012.

[23] Bahman Tahayori, Hamish Meffin, Socrates Dokos, Anthony N Burkitt, and David B Grayden. Modeling extracellular electrical stimulation: Ii. computational validation and numerical results. J. Neural Eng., 9(6):065006, 2012.

[24] Costas A Anastassiou and Christof Koch. Ephaptic coupling to endogenous electric field activity: why bother? Current opinion in neurobiology, 31:95–103, 2015.

[25] Chia-Chu Chiang, Rajat S Shivacharan, Xile Wei, Luis E Gonzalez-Reyes, and Dominique M Durand. Slow periodic activity in the longitudinal hippocampal slice can self-propagate non-synaptically by a mechanism consistent with ephaptic coupling. The Journal of physiology, 597(1):249–269, 2019.

[26] Andreas Fellner, Amirreza Heshmat, Paul Werginz, and Frank Rattay. A finite element method framework to model extracellular neural stimulation. Journal of Neural Engineering, 19(2):022001, 2022.

[27] Sébastien Joucla, Alain Gliére, and Blaise Yvert. Current approaches to model extracellular electrical neural microstimulation. Frontiers in computational neuroscience, 8:13, 2014.

[28] Wenjun Ying and Craig S Henriquez. Hybrid finite element method for describing the electrical response of biological cells to applied fields. IEEE. Trans. Biomed. Eng., 54(4):611–620, 2007.

[29] Andres Agudelo-Toro and Andreas Neef. Computationally efficient simulation of electrical activity at cell membranes interacting with self-generated and externally imposed electric fields. J. Neural Eng., 10:026019–026019, 2013.

[30] Leslie Tung. A bi-domain model for describing ischemic myocardial dc potentials. PhD thesis, Massachusetts Institute of Technology, 1978.

[31] Bradley J Roth. How the anisotropy of the intracellular and extracellular conductivities influences stimulation of cardiac muscle. Journal of mathematical biology, 30:633–646, 1992.

[32] ACL Barnard, IM Duck, and MS Lynn. The application of electromagnetic theory to electrocardiology: I. derivation of the integral equations. Biophys. J., 7(5):443–462, 1967.

[33] David B Geselowitz. On bioelectric potentials in an inhomogeneous volume conductor. Biophys. J., 7(1):1–11, 1967.

[34] Maurice Klee and Robert Plonsey. Integral equation solution for biopotentials of single cells. Biophysical Journal, 12(12):1676–1685, 1972.

[35] Luis J Gomez, Moritz Dannhauer, Lari M Koponen, and Angel V Peterchev. Conditions for numerically accurate tms electric field simulation. Brain Stimul., 13:157–166, 2020.

[36] Sergey N Makarov, Gregory M Noetscher, and Ara Nazarian. Low-frequency electromagnetic modeling for electrical and biological systems using MATLAB. John Wiley & Sons, 2015.

[37] Aung Thu Htet, Guilherme B Saturnino, Edward H Burnham, Gregory M Noetscher, Aapo Nummenmaa, and Sergey N Makarov. Comparative performance of the finite element method and the boundary element fast multipole method for problems mimicking transcranial magnetic stimulation (tms). J. Neural Eng., 16(2):024001, 2019.

[38] A L Hodgkin and A F Huxley. A quantitative description of membrane current and its application to conduction and excitation in nerve. 1952. Bull. Math. Biol., 52:25–71; discussion 5–23, 1952.

[39] J. A. Connor and C. F. Stevens. Inward and delayed outward membrane currents in isolated neural somata under voltage clamp. The J. Physiol., 213:1–19, 1971.

[40] P. Dayan and L.F. Abbott. Theoretical Neuroscience: Computational and Mathematical Modeling of Neural Systems. Computational Neuroscience Series. Massachusetts Institute of Technology Press, 2001.

[41] AG Richardson, CC McIntyre, and WM Grill. Modelling the effects of electric fields on nerve fibres: influence of the myelin sheath. Medical and Biological Engineering and Computing, 38:438–446, 2000.

[42] John Crank and Phyllis Nicolson. A practical method for numerical evaluation of solutions of partial differential equations of the heat-conduction type, volume 43. 1947.

[43] J. W. (James William) Thomas. Numerical partial differential equations : finite difference methods. Texts in applied mathematics ; 22. Springer, New York, 1st ed. 1995. edition, 1995.

[44] Olaf Steinbach. Numerical approximation methods for elliptic boundary value problems: finite and boundary elements. Springer Science & Business Media, 2007.

[45] Tianyi Zhou, Yixuan Ming, Susan F Perry, and Svetlana Tatic-Lucic. Estimation of the physical properties of neurons and glial cells using dielectrophoresis crossover frequency. Journal of biological physics, 42:571–586, 2016.

[46] Zhi-De Deng, Sarah H Lisanby, and Angel V Peterchev. Electric field depth–focality tradeoff in transcranial magnetic stimulation: simulation comparison of 50 coil designs. Brain stimulation, 6(1):1–13, 2013.

[47] Luis Gomez, Frantishek Cajko, Luis Hernandez-Garcia, Anthony Grbic, and Eric Michielssen. Numerical analysis and design of single-source multicoil tms for deep and focused brain stimulation. IEEE. Trans. Biomed. Eng., 60:2771–2782, 2013.

[48] Luis J Gomez, Abdulkadir C Yucel, Luis Hernandez-Garcia, Stephan F Taylor, and Eric Michielssen. Uncertainty quantification in transcranial magnetic stimulation via high-dimensional model representation. IEEE. Trans. Biomed. Eng., 62:361–372, 2015.

[49] P J Maccabee, V E Amassian, L P Eberle, and R Q Cracco. Magnetic coil stimulation of straight and bent amphibian and mammalian peripheral nerve in vitro: locus of excitation. The J. Physiol., 460:201–219, 1993.

[50] Gary R Holt and Christof Koch. Electrical interactions via the extracellular potential near cell bodies. Journal of computational neuroscience, 6:169–184, 1999.

[51] Aaron R Shifman and John E Lewis. Elfenn: a generalized platform for modeling ephaptic coupling in spiking neuron models. Frontiers in Neuroinformatics, 13:35, 2019.

[52] Yamina Bakiri, Ragnhildur Káradóttir, Lee Cossell, and David Attwell. Morphological and electrical properties of oligodendrocytes in the white matter of the corpus callosum and cerebellum. The Journal of physiology, 589(3):559–573, 2011.

[53] Wanda Krassowska and John C Neu. Effective boundary conditions for syncytial tissues. IEEE transactions on biomedical engineering, 41(2):143–150, 1994.

[54] Hemant Bokil, Nora Laaris, Karen Blinder, Mathew Ennis, and Asaf Keller. Ephaptic interactions in the mammalian olfactory system. The Journal of neuroscience, 21(20):RC173, 2001.

[55] Ada J Ellingsrud, Andreas Solbrå, Gaute T Einevoll, Geir Halnes, and Marie E Rognes. Finite element simulation of ionic electrodiffusion in cellular geometries. Frontiers in Neuroinformatics, 14:11, 2020.

[56] Abtin Rahimian, Ilya Lashuk, Shravan Veerapaneni, Aparna Chandramowlishwaran, Dhairya Malhotra, Logan Moon, Rahul Sampath, Aashay Shringarpure, Jeffrey Vetter, Richard Vuduc, Denis Zorin, and George Biros. Petascale direct numerical simulation of blood flow on 200k cores and heterogeneous architectures. In SC ‘10: Proceedings of the 2010 ACM/IEEE International Conference for High Performance Computing, Networking, Storage and Analysis, pages 1–11, 2010.

[57] David M. Czerwonky and Luis J. Gomez. Integral equation for analyzing neuron response to non-invasive electromagnetic brain stimulation. In 2023 International Applied Computational Electromagnetics Society Symposium (ACES), pages 1–2, 2023.

[58] Robert Plonsey. The formulation of bioelectric source-field relationships in terms of surface discontinuities. Journal of the Franklin Institute, 297(5):317–324, 1974.

