## Supplementary material for "A Boundary Element Method of Bidomain Modeling for Predicting Cellular Responses to Electromagnetic Fields": Bidomain BEM Supplemental

**Address:** Seng-Liang Wang Hall Rm 3059, 516 Northwestern Ave, West Lafayette, IN 47906

### Bidomain BEM Supplementary Materials

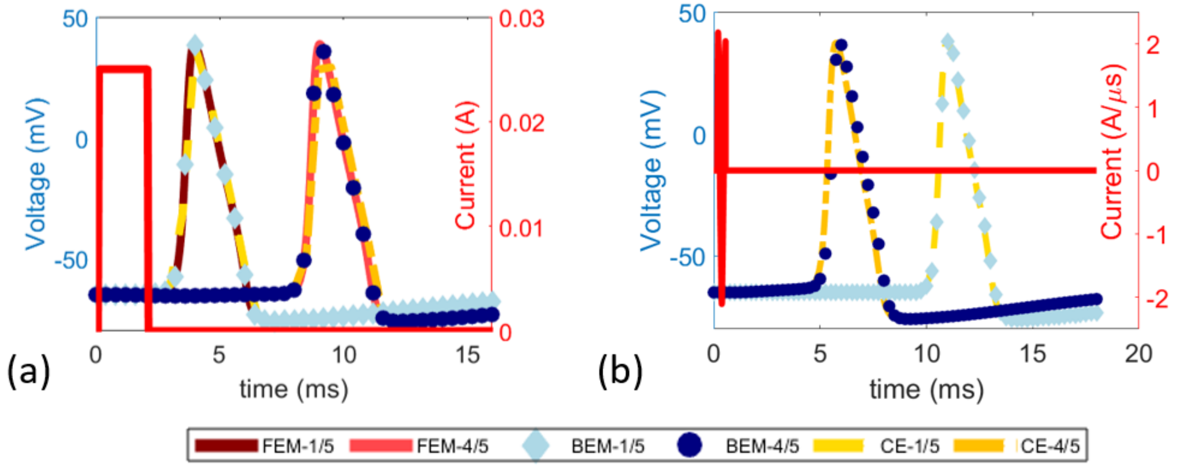


**Figure S1**: Supplemental data of the longitudinal comparison scenarios. (a) TES results comparing BEM, FEM, and the cable equation (CE). (b) TMS results comparing BEM and the cable equation (CE). We show here that our bidomain BEM can perform the same as its FEM equivalent in addition to its performance with respect to the cable equation as shown in the main publication. There is no bidomain FEM platform available for TMS, so we did not perform any additional comparisons with TMS.

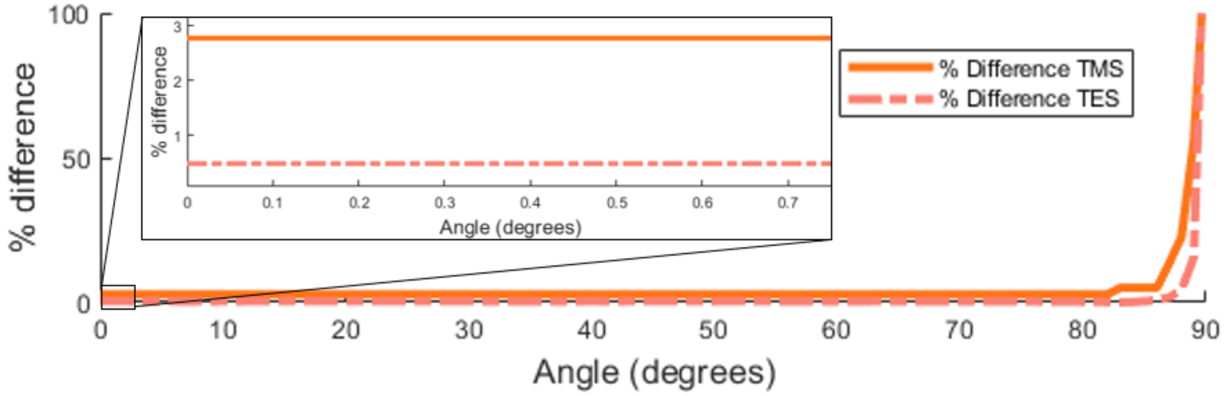


**Figure S2**: Supplemental data of percent differences between the bidomain BEM and the cable equation from the axon rotation scenarios for TES and TMS (figures 7 and 8 of the main publication), respectively. The difference is less than 3 everywhere under 83° for TMS and 89° for TES. At 90°, the difference becomes extremely large since the cable equation cannot predict transverse stimulation.

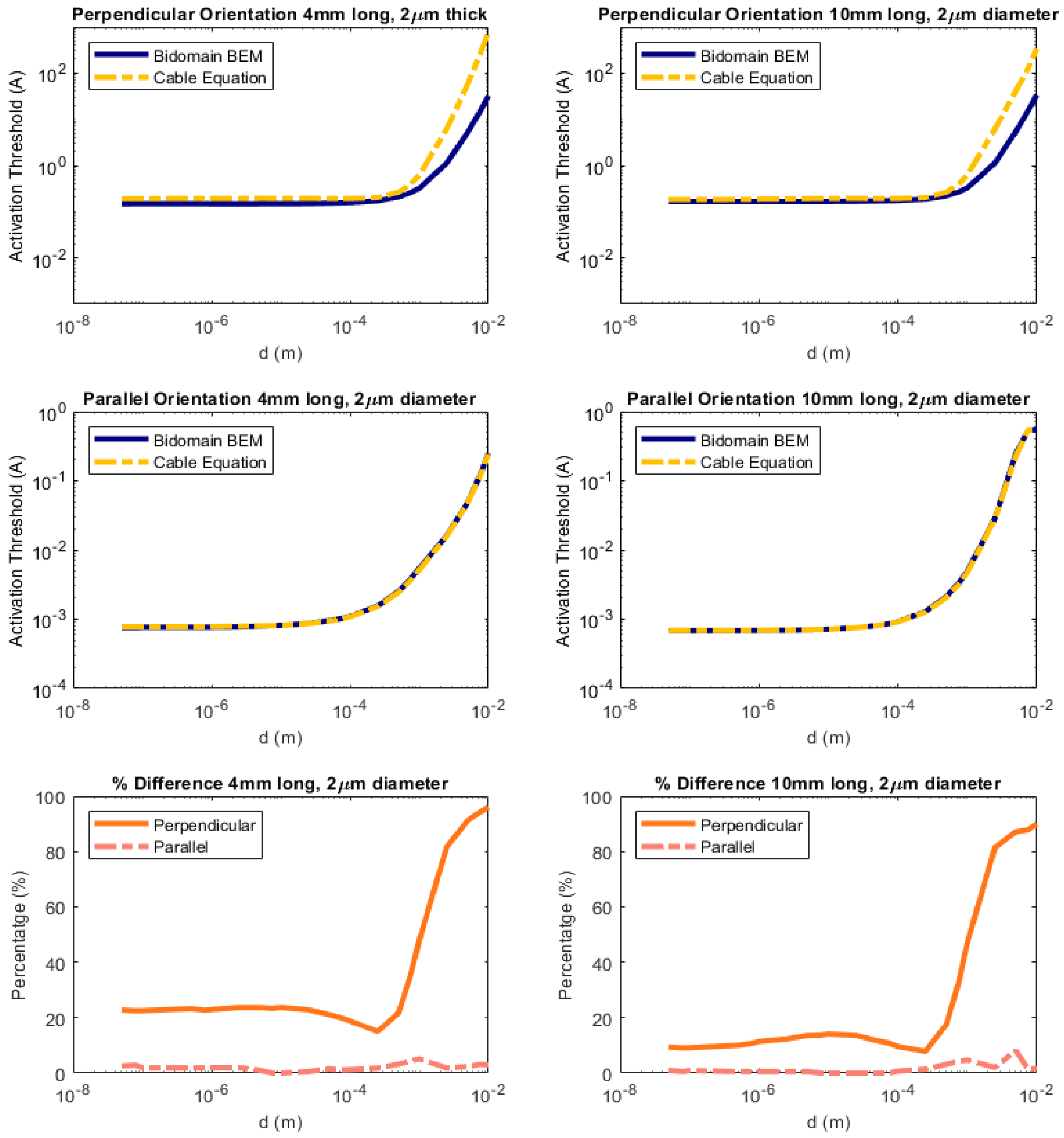


**Figure S3**: (Row 1): Plots of the activation threshold for different axon geometries perpendicular to a DBS electrode at a distance d from the axon. (Row 2): Plots of the activation threshold for different axon geometries parallel to a DBS electrode at a distance d from the axon. (Row 3) Plots of the percentage difference between bidomain BEM and the cable equation for each respective geometry.

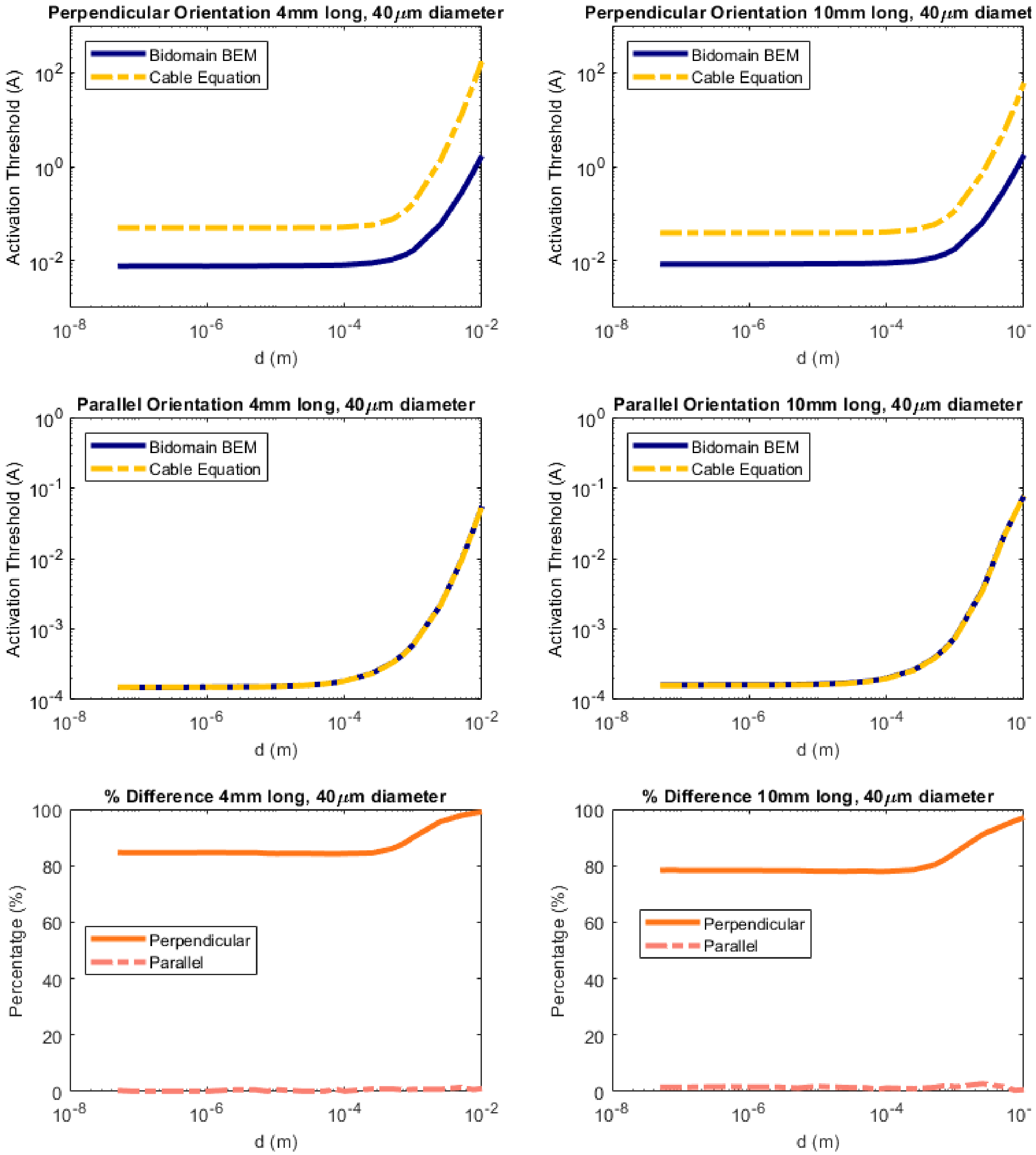


**Figure S4**: (Row 1): Plots of the activation threshold for different axon geometries perpendicular to a DBS electrode at a distance d from the axon. (Row 2): Plots of the activation threshold for different axon geometries parallel to a DBS electrode at a distance d from the axon. (Row 3) Plots of the percentage difference between bidomain BEM and the cable equation for each respective geometry.

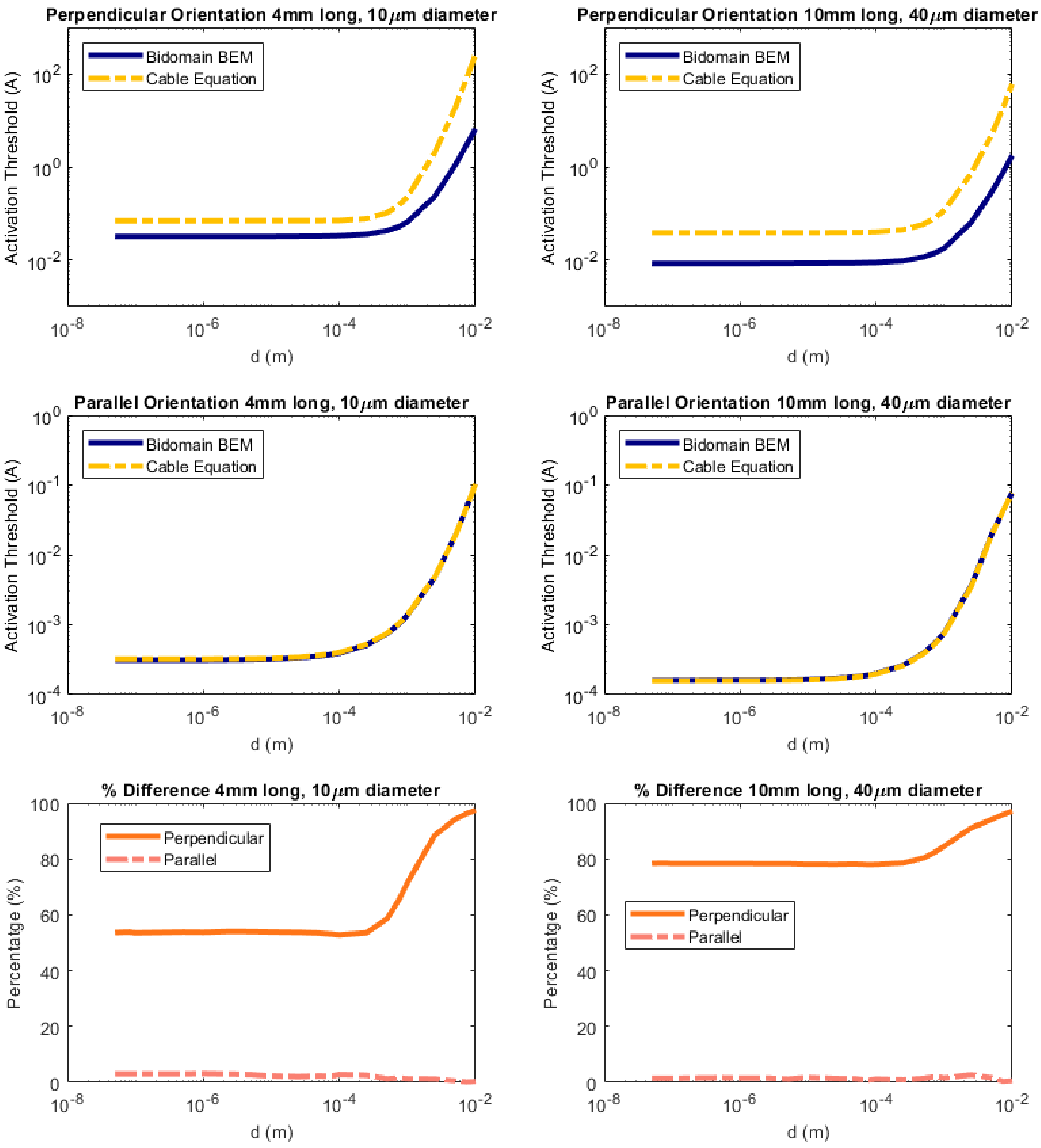


**Figure S5**: (Row 1): Plots of the activation threshold for different axon geometries perpendicular to a DBS electrode at a distance d from the axon. (Row 2): Plots of the activation threshold for different axon geometries parallel to a DBS electrode at a distance d from the axon. (Row 3) Plots of the percentage difference between bidomain BEM and the cable equation for each respective geometry.

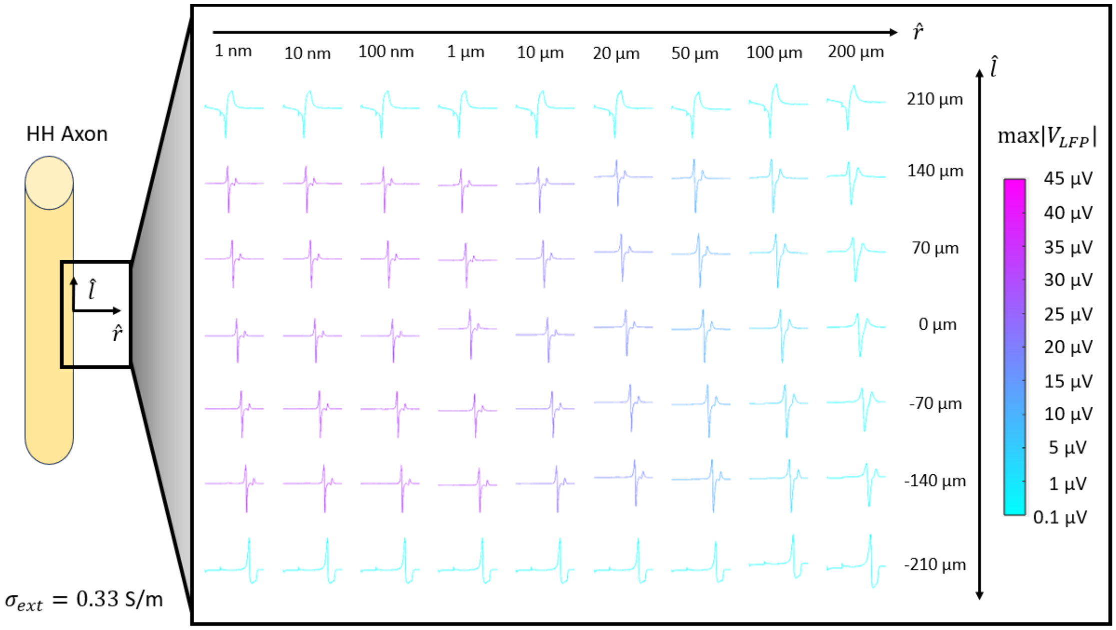


**Figure S6**: Supplemental data of local field potentials at a higher conductivity value in a realistic range than given in the main article. A schematic detailing the region where the data originates is on the left. The plot shows local field potentials waveforms sampled at the positions on the l-r grid. The colors indicate the magnitude of the largest peak of each respective voltage waveform.

### Bidomain BEM Supplementary Materials

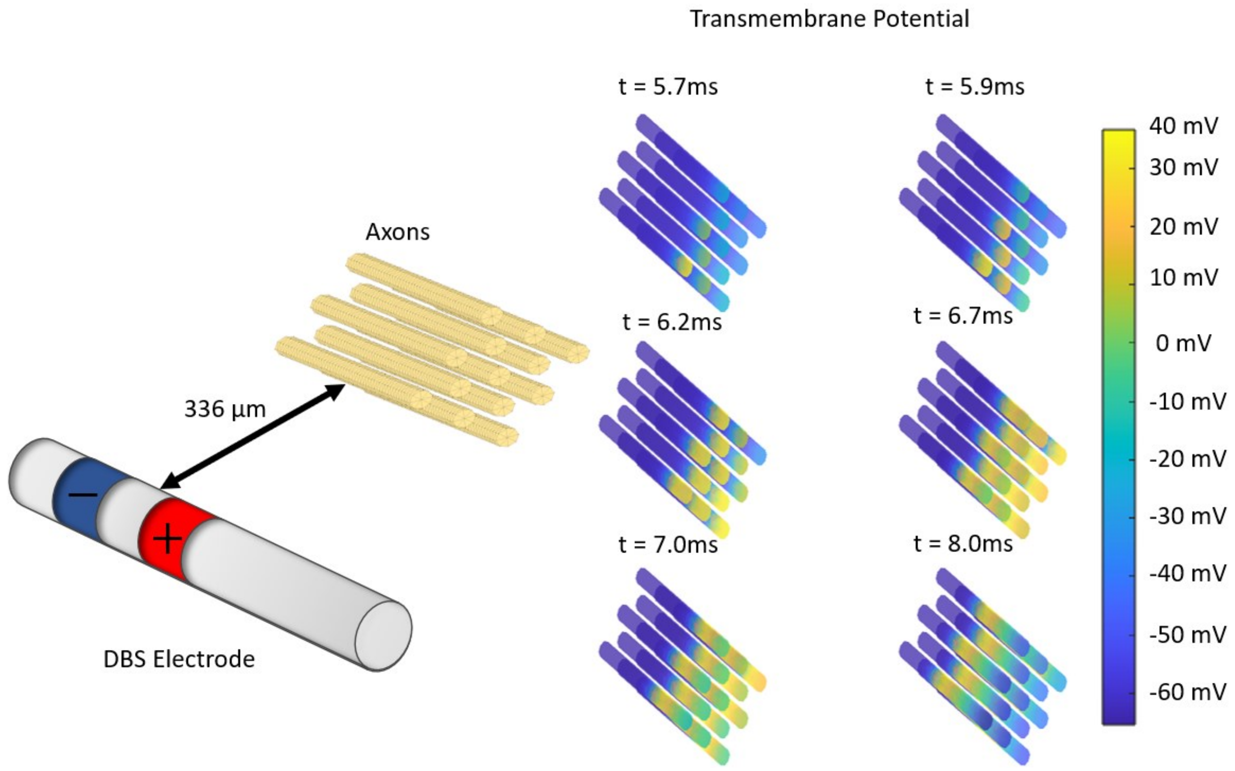


**Figure S7**: Supplemental data of device stimulation of a bundle of 13 axons with 1.4 μm spacing between them. The DBS electrode was driven with a titrated current of 2 mA. This activated all the cells in a cascade as shown above. The cascade is unrelated to ephaptic interactions because the same effects were observed with each axon in isolation.

| Cell ID | Isolated | Conductors | Active Cells | Insulators |
|---|---|---|---|---|
| 1 | 5.36 mA | 5.36 mA | 6.20 mA | 6.20 mA |
| 2 | 5.36 mA | 4.83 mA | 5.89 mA | 5.89 mA |
| 3 | 5.36 mA | 5.56 mA | 5.52 mA | 5.53 mA |
| 4 | 5.36 mA | 5.22 mA | 5.89 mA | 5.88 mA |
| 5 | 5.36 mA | 4.93 mA | 5.90 mA | 5.90 mA |
| 6 | 5.36 mA | 5.22 mA | 5.89 mA | 5.88 mA |
| 7 | 5.36 mA | 5.56 mA | 5.53 mA | 5.53 mA |

(A)

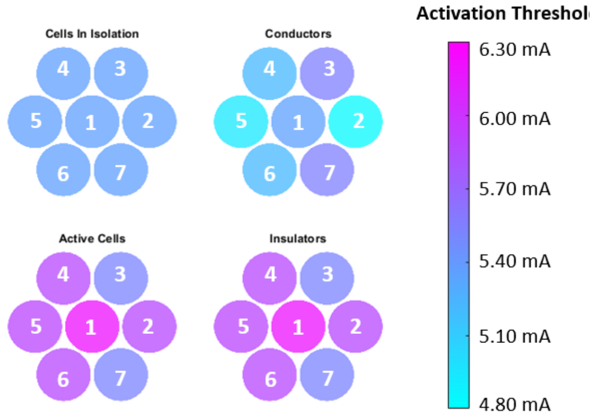


(B)

**Figure S8:** Table of activation thresholds in a 0.1 S/m extracellular space that compares individual cells (isolated) to a network of all seven cells in a hexagonal configuration while all cells, but the cell of interest are passive, active, or under initial polarization conditions i.e. an insulator. (B) Color plots of the threshold values with respect to each soma and the properties of its surrounding cells.

### Bidomain BEM Supplementary Materials

| Cell ID | Isolated | Conductors | Active Cells | Insulators |
|---|---|---|---|---|
| 1 | 107 mA | 111 mA | 124 mA | 124 mA |
| 2 | 107 mA | 111 mA | 118 mA | 118 mA |
| 3 | 107 mA | 108 mA | 110 mA | 110 mA |
| 4 | 107 mA | 110 mA | 118 mA | 118 mA |
| 5 | 107 mA | 111 mA | 118 mA | 118 mA |
| 6 | 107 mA | 110 mA | 118 mA | 118 mA |
| 7 | 107 mA | 108 mA | 110 mA | 110 mA |

**Table S1:** Table of activation thresholds in a 0.1 S/m extracellular space that compares individual cells (isolated) to a network of all seven cells in a hexagonal configuration while all cells, but the cell of interest are passive, active, or under initial polarization conditions i.e. an insulator.
